# Fabrication of Short Polymeric µFibers as Building Blocks for Anisotropic High-Throughput Compatible 3D Tissue Models

**DOI:** 10.64898/2025.12.09.692980

**Authors:** Anna A. Meyer, Abdolrahman Omidinia-Anarkoli, Michelle Harmeth, Joscha Graeve, Karoline Rengel, Melike Keskin, Ramin Nasehi, Maximilian Fusenig, Nicholas R. Dennison, Till Hülsmann, Tamás Haraszti, Uwe Freudenberg, Carsten Werner, Laura De Laporte

## Abstract

Modeling the 3D microenvironment and cell growth of natively anisotropic human tissues *in vitro* constitutes a significant challenge in tissue engineering and biofabrication. Short polymeric fibers gain growing attention in this field due to their applicability as pipettable or injectable anisometric building blocks in 3D hydrogel-based cell culture systems or bioinks. However, the production of monodisperse short fibers with high production rates suitable for screening remains challenging. In this study, short, quasi-monodisperse, magneto-responsive, fluorescent poly(ε-caprolactone) µfibers with variable dimensions in the micrometer range are produced in a scalable, semi-continuous two-step fabrication process combining controlled wet-dry spinning with subsequent cryosectioning. Influences of the spinning process parameters on fiber properties and process features, as well as boundary spinning conditions and upscaling potential, are explored using Design of Experiments approaches. Further, magnetic alignment of the µfibers in a weak magnetic field and incorporation of nile red as fluorescent dye for facile analysis in 3D are demonstrated. Implementation of aligned µfibers into a hydrogel-based 3D vasculogenesis model, produced in a high-throughput automated manner, is shown to stimulate oriented cell growth. This highlights the potential of our µfibers as guiding elements inside tissue and disease models and their suitability for automated high-throughput applications.

## 1 Introduction

As the production of three-dimensional (3D) tissue models that mimic the native structure and function of human tissues still constitutes a challenge in tissue engineering and regenerative medicine,^[1,2]^ there is an ongoing need for the development of novel biomaterials and building blocks. Over the last two decades, fiber-based scaffolds, enabling structural stability and increased porosity for nutrient diffusion, as well as mimicking the structural properties of native extracellular matrix (ECM), gained growing attention.^[3,4,5]^ Poly(ε-caprolactone) (PCL) is a polymer widely used to manufacture polymeric fibers or fiber scaffolds due to its versatile processability, biocompatibility and slow degradation rate, and therefore has been extensively used for medical devices.^[6–8]^ It can be processed in melt, wet, dry or electrospinning applications.^[6,9–11]^ Electrospinning is the most common technique to fabricate PCL fibers and yields fiber diameters in the nano- to micrometer range,^[4,7]^ which match the diameter of natural ECM fibers.^[5,12]^ In contrast, wet and dry spinning typically yield thicker fibers with diameters in the range of tens to hundreds of micrometers,^[9,13]^ although, depending on specific setup and drawing ratios, fiber diameters below 10 µm were successfully produced with dry spinning.^[14]^ As fiber spinning is governed by several process parameters like flow rate, drum speed, distance between cannula and collector and voltage (in electrospinning), Design of Experiments (DoE) approaches, employing statistical experimental planning, are implemented more frequently to characterize fiber spinning processes.^[9,15,16,17]^ DoE enables efficient experimental work by yielding the highest amount of information with the lowest number of experimental runs possible.^[18]^ It is used to systematically analyze effects of process parameters, detect possible parameter interactions and establish prediction models for investigated properties based on polynomial regression analysis.^[19]^

In addition to the applications of fibrous scaffolds mentioned above, fiber-based materials were successfully employed over the past decade to mimic anisotropic tissues like nerves, heart, tendon or muscle tissue, exploiting the anisotropic nature of fibers.^[2,20]^ Besides aligned fiber mats^[5,21]^ or scaffolds with controlled inter-fiber angles,^[15,22]^ fragmented short fibers with lengths in the micrometer range were developed as anisometric building blocks for 3D cell culture models.^[23–28]^ In contrast to large fibrous scaffolds, short fibers are pipettable and injectable, thereby permitting facile incorporation in widely used hydrogel-based *in vitro* cell culture models, as well as applications in minimally invasive therapies or 3D bioprinting. Although even randomly oriented short fibers co-embedded with cells in a hydrogel matrix could enhance cell differentiation and oriented cell growth,^[24]^ alignment of short fibers along one direction was shown to be a powerful tool to trigger highly anisotropic cell growth.^[20]^ In this regard, the Anisogel technology was developed by our group, enabling magnetic alignment of magneto-responsive electrospun short fibers dispersed in a hydrogel precursor solution by applying a weak, external magnetic field; crosslinking of the surrounding hydrogel fixes the orientation of the rods, thereby allowing the removal of the magnetic field after gelation.^[23,29]^ Here, it is important that the fibers align before the hydrogel is crosslinked, thus the kinetics of both processes have to conform to each other.

Fabrication of short fibers can be achieved either by in-situ fragmentation of fibers during their production process (e.g. electric spark-assisted electrospinning,^[30]^ entropy-induced fiber snapping during electrospinning,^[31]^ shear precipitation of polymer solution injected in non-solvent^[27,32]^) or, as more commonly done, by spinning of continuous fibers and subsequent fiber fragmentation.^[33,34]^ Post-spinning fragmentation relies on chemical or mechanical fiber scission techniques. While the applicability of chemical scission techniques (e.g. aminolysis,^[35]^ UV crosslinking with patterned photomasks^[36]^) strongly depends on the fiber material, mechanical scission can be used in a more universal manner for a broad range of polymeric or hydrogel-based fiber materials. Scission by repeated ejection of fiber mats through a needle with subsequent filtering,^[24,37]^ milling and sieving,^[27]^ ultrasonication,^[38]^ homogenization with a blender or rotating razor blades,^[25,39]^ and cryosectioning with a microtome^[23,26,28,40,41,42]^ have been presented in literature for mechanical fiber fragmentation, resulting in fiber lengths in the range of a few to several hundred micrometers.^[33]^ Here, cryosectioning offers the most facile method regarding the control of resulting fiber length as the parameter can be adjusted directly via the microtome cutting thickness, while, in the other methods, the fiber length is indirectly affected by multiple different parameters, e.g. repetition cycles or duration of the applied technique, dispersion medium, frequencies or rotation speeds. Despite this, cryosectioning often yields imprecise fiber lengths and broad length distributions, for example an average fiber length of 69 µm for 50 µm thick sectioning slices and standard deviations ranging from 25 to 70% of the average fiber length.^[26,40,42]^ The sectioning precision can be enhanced by improving the degree of alignment of spun fibers prior to sectioning. However, aligned fiber collection during electrospinning, the spinning method used almost exclusively in literature to produce short fibers, is challenging due to the typical whipping motion of the fiber between nozzle and collector. Although fiber alignment during collection can be controlled to some extent by using rotating collectors at high rotation speeds^[43]^ or a careful selection of process parameters,^[17]^ high degrees of alignment require more elaborate electrospinning setups^[43,44]^ so far not commonly used for the fabrication of short fibers.

Alternatively, fabrication of more monodisperse polymeric^[45]^ or hydrogel-based^[46,47,48]^ micrometer-sized building blocks for anisotropic 3D cell culture models is possible (using e.g. micropatterning and spin-coating techniques, particle replication in non-wetting templates (PRINT), or jet microfluidics) but usually entails time- and/or labor-intensive batch processes, making upscaling challenging. Especially when targeting the production of high-throughput (HT) tissue models, e.g., for drug screening to reduce animal testing and improve personalized medicine,^[2]^ respective building blocks for these models need to be fabricated in large scale.

In this study, we present a scalable, semi-continuous fabrication method of magneto-responsive short PCL fibers (µfibers) that can be used as anisotropic building blocks in HT 3D tissue and disease models (Figure 1, top row). The fibers, featuring diameters in the low micrometer range, are produced in a wet-dry spinning process allowing fiber collection in a highly aligned manner on a rotating drum (due to the absence of the in electrospinning typically observed whipping motion of the fiber), and are subsequently cryosectioned into short µfibers with precise lengths. The influence of spinning process parameters on fiber diameter and other fiber properties and process features is systematically investigated using DoE. Based on polynomial regression analysis of the empirical data, a correlative model to predict the fiber properties and process features, also at untested parameter combinations, is obtained. The upscaling potential of the spinning process is explored and alignment of the µfibers in an external magnetic field and µfiber fluorescence are analyzed. To demonstrate the relevance of the µfibers as building blocks for HT tissue models, µfibers are implemented into a hydrogel-based co-culture of human umbilical vein endothelial cells (HUVECs) and human mesenchymal stem cells (MSCs) (schematically displayed in Figure 1, bottom row). Using an automated fluid handling system and cell culture workstation for preparation of the 3D cell culture models allowed for a systematic study of the influence of the aligned µfibers on the orientation of vascular sprouting and proved the HT compatibility of this technology.

**Figure 1:**
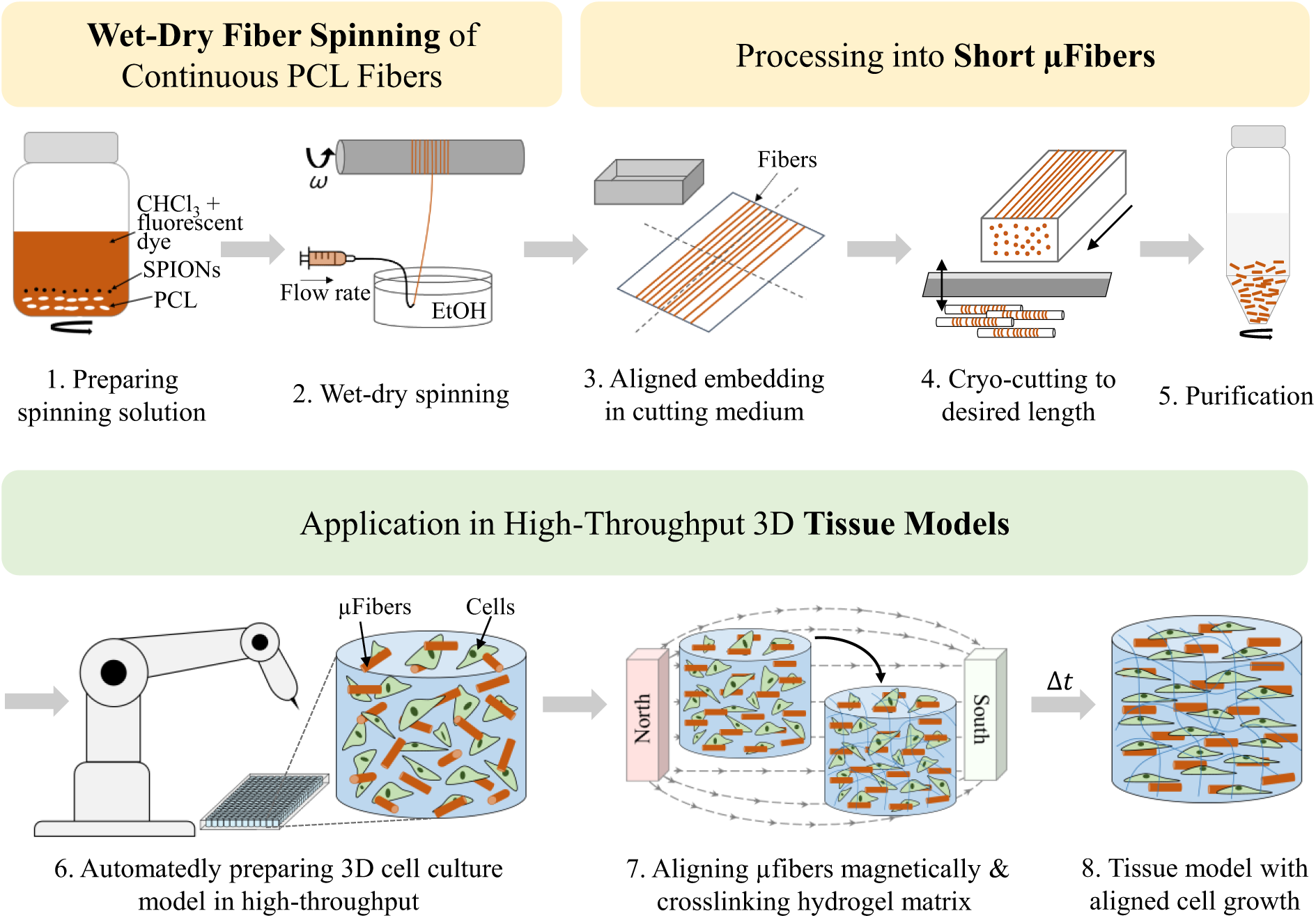
Schematic presentation of the production process of short, magneto-responsive µfibers and their possible application in high-throughput 3D tissue models. The µfiber production process includes the following steps: (1) preparation of the spinning solution, (2) wet-dry spinning, (3) aligned embedding of spun fibers into a cutting medium, (4) cryosectioning to a desired µfiber length, (5) purification of the µfibers (and disinfection if further used in cell culture). The µfibers can be used as anisometric building blocks in 3D tissue models by: (6) preparing 3D cell culture models with µfibers in an automated manner using a fluid handling system, (7) magnetically aligning the µfibers with an external magnetic field (possible directly within the fluid handling system) and crosslinking the surrounding hydrogel matrix. µFibers will function as anisometric guiding elements during incubation, yielding (8) a 3D tissue model with aligned cell growth.

## 2 Results and Discussion

### 2.1 Development of µFiber Production Process and Wet-Dry Spinning Setup

For production of polymeric magneto-responsive micrometer-sized rods as guiding elements for 3D tissue culture in high quantity and a semi-continuous manner, we developed a wet-dry spinning process with subsequent cryosectioning (Figure 1, top row). To prepare the spinning solution (Figure 1 (1)), PCL is dissolved in chloroform. Superparamagnetic iron oxide nanoparticles (SPIONs) are added to render the resulting µfibers magneto-responsive and thus enable an alignment along a weak magnetic field (∼ 10 − 40 mT) for future applications in tissue culture. Further, a fluorescent dye is added to enable analysis of the µfibers with confocal fluorescence microscopy in 3D hydrogel constructs. As we selected a combination of wet and dry spinning for our process (Figure 1 (2)), lower flow rates and the absence of an electric field (i.e. no whipping motion of the filament) allow collection of the fibers in a more controlled and aligned manner on a rotating collector compared to commonly used electrospinning processes. The aligned collection of fibers is crucial for obtaining of monodisperse fiber fragments during subsequent cryosectioning. After spinning, the fibers are harvested from the drum, embedded in an optimal cutting temperature medium (OCT) in an aligned manner (Figure 1 (3)) and sectioned into µfibers with defined lengths (Figure 1 (4)). For this study, we chose 50 µm as standard µfiber length based on previous studies on the influence of rod-shaped elements used as guiding elements in cell culture.^[47]^ µFibers are purified by washing with water to remove the water-soluble cryosectioning medium (Figure 1 (5)). Analyzing the wash supernatant by lyophilization and subsequent infrared spectroscopy revealed that at least four washing cycles are necessary to remove the OCT from the µfiber dispersion (data not shown).

In the µfiber production process, the wet-dry spinning is a crucial step as it determines the µfiber diameter and morphology as well as the production rate of µfibers through the stability of the spinning process. The spinning setup is comprised of a syringe pump for extrusion of the spinning solution through a customized cannula pointing upwards, a coagulation bath and a rotating drum (coated with dried OCT cryosectioning medium) positioned above. For fiber collection, spinning solution extruded into the coagulation bath is picked up as a coagulating filament and placed onto the rotating drum (Figure 1 (2)). Through coagulation and subsequent drying in the air gap between coagulation bath and collector, a filament forms which is continuously collected on the drum.

Process features and fiber properties are influenced by multiple material and process parameters and environmental conditions, such as the molecular weight and concentration of the polymer in the spinning solution, coagulation bath solvent, cannula diameter, flow rate, drum speed, distance between drum and cannula (vertically and horizontally), temperature and humidity. First, varying PCL concentrations were tested under similar spinning conditions (flow rate: 0.3 mL/h, drum speed: 78.5 cm/s, similar positioning of cannula and drum) to investigate which range of polymer concentrations in our spinning solution allows for stable spinning. Concentrations between 10 and 25 wt/v% were found to be spinnable, whereas lower or higher concentrations were either too fluid or too viscous to form a stable filament in the coagulation bath. Resulting fiber diameters, measured based on microscopy images, increased with increasing concentration up to 15 wt/v%; above this concentration, the diameter reached a plateau (Figure S1, Supporting Information). For following spinning experiments of this study, a PCL concentration of 17 wt/v%, yielding fiber diameters within the observed plateau and being comparable to previous studies,^[23,46]^ was used.

As solvent for the coagulation bath, ethanol is employed as it is a non-solvent for PCL, enhancing fiber solidification through a phase inversion induced by solvent/non-solvent exchange.^[11,49]^ The coagulation bath is necessary for our spinning process to enable a continuous fiber spinning. Without a coagulation bath but otherwise same setup, i.e. in a dry spinning process, the extruded filament rips off while pulling it towards the drum or, if it could be placed on the drum, after a few seconds of spinning. The chloroform evaporates very quickly after the spinning solution exits the cannula, making the filament brittle and prone to clogging the cannula tip during dry spinning. Similarly, no continuous dry spinning could be achieved by placing the extrusion nozzle above the rotating collector and extruding along the gravity force direction to produce fibers in our targeted low micrometer range. Previous reports employed dry spinning of PCL dissolved in dichloromethane using a larger nozzle diameter; however, this method produced fibers with diameters in the range of several hundred micrometers.^[9]^ To investigate the effect of the coagulation bath on the fiber diameter, the standard spinning solution was extruded into the ethanol bath without subsequent collection on the drum to avoid any stretching of the filament. Resulting fibers exhibited a fiber diameter of 90.7 ± 9.2 µm, indicating that the filament shrinks to circa 17% of the inner cannula diameter (0.514 mm for 21G cannula) through coagulation. Further reduction of the diameter of fibers produced in this study can therefore be attributed to stretching by collecting the fibers on the rotating drum.

During development of the spinning setup, two preliminary stages of the setup were tested before the final setup, used for further experiments, was obtained (Figure S2, Supporting Information). Starting from the first preliminary setup, the centering of the drum was enhanced by fixing the drum to its rotational axis on both sides to improve the stability of the spinning process, i.e. increase the time of continuous spinning without the fiber ripping off the drum. Thereby, fiber movement on the drum was decreased because of smoother drum rotation, enabling continuous spinning of up to 8 h (stopped manually then because work day ended). Further, a precise drum speed control based on a stepper motor was implemented and the drum holder was mounted onto a rail allowing stepless drum height variation for the investigation of the influence of these process parameters on the spinning process and resulting µfiber properties. With the final spinning setup (Figure S2c, Supporting Information), PCL fibers with a smooth surface and cross-section morphology and diameters in the low micrometer range (Figure 2a and b) are obtained; with subsequent cryosectioning, as described above, yielding µfibers with a defined length (Figure 2c). For instance, sectioning of fibers with slice thicknesses of 25, 50 or 100 µm programmed at the microtome resulted in actual µfiber lengths of 27 ± 1.9, 55 ± 3.5, or 108 ± 5.9 µm, respectively; the obtained lengths thus match the targeted lengths more closely and standard deviations of 5 to 7% of the average fiber length indicate significantly narrower length distributions than respective values found in literature.^[26,40]^ µFibers featuring these narrower length distributions constitute desirable building blocks for tissue engineering purposes as they allow for studying the effect of different building block lengths on cell behavior with enhanced precision.

**Figure 2:**
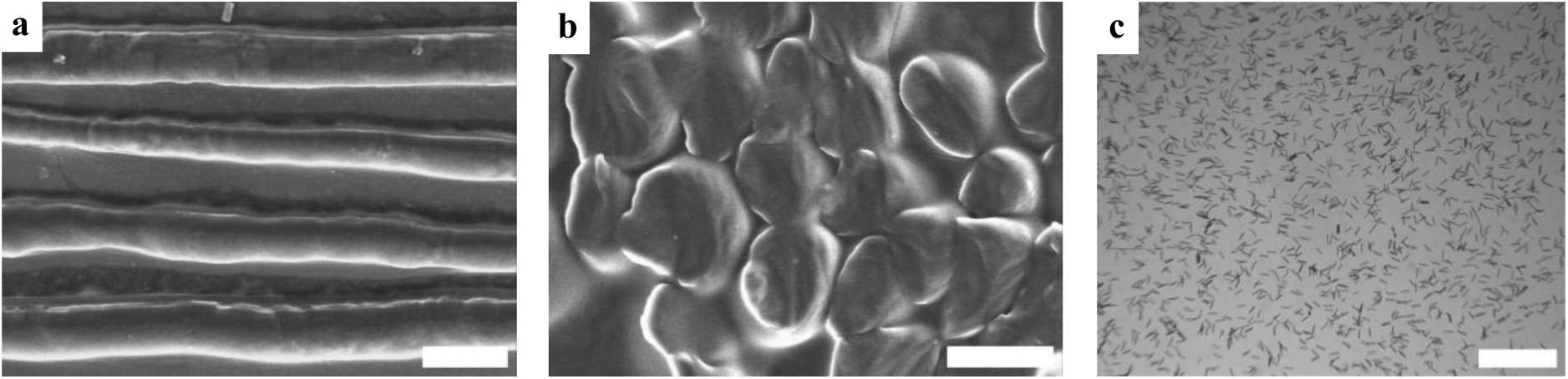
Scanning electron microscopy (SEM) images of (a) fiber surface and (b) fiber cross-sections. Scale bars: (a) 10 µm, and (b) 5 µm. (c) Microscopy image of µfibers cut to 50 µm length, dispersed in water. Scale bar: 500 µm.

### 2.2 Influence of Process Parameters on Fiber Properties and Process Features

For efficient large-scale production of µfibers with specific dimensions, a stable spinning process and a thorough understanding of the process parameters’ influence on fiber properties and process features is essential. Based on requirements for a robust large-scale fabrication of µfibers, fiber diameter, µfiber merging, the average amount of fiber rip-offs per hour and the spinning area of the fibers on the drum after spinning were identified as output parameters of interest (Figure 3a), which vary based on experimental settings during spinning. Targeted µfiber diameters for 3D cell alignment are in the range of micrometers^[47]^ and can be controlled during fiber spinning by applied flow rate and drum speed. Unwanted µfiber merging, i.e. two or more µfibers sticking together longitudinally forming inseparable fiber stacks, can be observed after µfiber cutting and purification. This is caused by insufficient solvent evaporation from the extruded filament before touching already collected fibers on the drum, thereby solvating the surface of the already collected fibers again and allowing polymer chains of both fibers to entangle before drying. For our targeted application of using µfibers as building blocks in 3D tissue cultures, single µfibers with controlled dimensions are desired, thus, merging should be avoided. How often the fiber rips off from the drum during spinning and the spinning area on the drum are process features relevant for possible upscaling of the process; continuous spinning without the fiber ripping off as well as a narrow spinning area on the drum are desirable for achieving fiber alignment and enabling a high throughput, parallelization of the process and efficient purification as less OCT is required to embed and cut the fibers.

**Figure 3:**
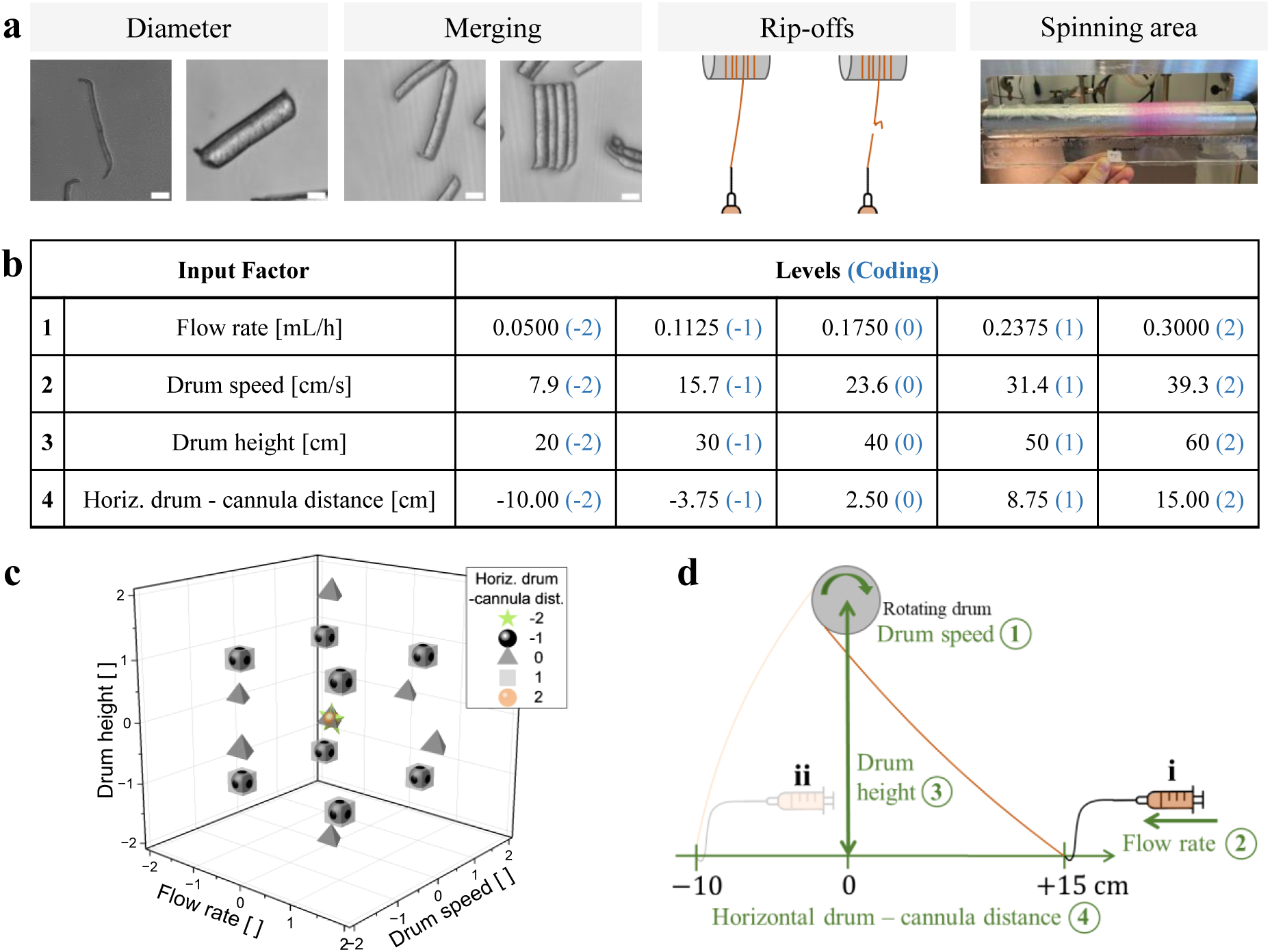
(a) Output parameters of interest for a robust production of µfibers in large quantaties; images show exemplary outputs. Scale bars: 10 µm. (b) To investigate their influence on the identified output parameters, the listed process parameters (i.e. input factors) were tested at different levels given in the table in a Design of Experiments approach (DoE I). Blue numbers in brackets give the values normalized to the middle value of the parameter range chosen for the respective input factor, referred to as coding in the following. (c) Combination of input factor levels tested in different DoE runs, selected based on a central composite design with star points (*α*_CCD_ = 2). The depicted levels refer to the coding introduced in (b). (d) Schematic presentation of the input factors (green) within the spinning setup: (1) drum speed of the rotating collector drum, (2) flow rate of the spinning solution, (3) drum height (vertical distance measured between center of drum and the horizontal plane where the cannula is positioned), (4) horizontal drum – cannula distance (horizontal distance of the cannula to position 0 below the drum). Positioning of the syringe shown for the two extreme settings of the horizontal drum – cannula distance at +15 cm (i) and −10 cm (ii).

In a first DoE approach (DoE I) employing a central composite design, the process parameters flow rate (0.05 − 0.3 mL/h), drum speed (37.5 − 187.5 rpm, with drum diameter of 4 cm resulting in surface speeds of 7.9 − 39.3 cm/s), drum height (vertical distance, 20 − 60 cm) and horizontal drum – cannula distance (−10 – 15 cm, i.e. cannula in side view left or right of clockwise rotating drum, as depicted in Figure 3d) were systematically varied as input factors (Figure 3b and c) to investigate their influence on the output parameters described above (Figure 3a). In total, 29 spinning experiments with different process parameter combinations, called DoE runs in the following, were conducted. Using polynomial regression analysis with standard least squares as fitting method on the experimental data (Figure S3, Supporting Information), the impact of the process parameters (main effects, e.g. *X*_1_, *X*_2_), as well as quadratic or non-linear parameter interactions (interaction effects, e.g. *X*^2^, *X*_1_*X*_2_), were explored and a prediction model for each output parameter was generated by computing second order polynomials for the respective output. Detailed accounts on the models, respective effect summary plots, prediction profilers and interaction plots can be found in the Supporting Information (Figure S4 – S8). Results for each output parameter as well as an experimental model validation are discussed in the following sections.

#### Fiber Diameter

In the experimental DoE runs, fiber diameters between 3.4 ± 0.7 µm (DoE run 3) and 12.0 ± 2.1 µm (DoE run 1) were obtained (Figure S3b, Supporting information). A negative linear effect of the drum speed was found to exert the most pronounced effect on the fiber diameter, closely followed by a positive linear effect of the flow rate, meaning that, with increasing drum speed, the fiber diameter will be decreased while increasing the flow rate will enlarge the fiber diameter. Besides their linear effects, quadratic influences appeared at higher drum speeds or flow rates. Furthermore, an interaction effect of drum speed and flow rate was observed as the fiber diameter is more affected by the flow rate at lower drum speeds (Figure 4a and b; and Figure S5, Supporting information). From a physical point of view, these trends were expected as a higher drum speed will lead to a higher degree of filament stretching and a thinner fiber whereas a higher flow rate results in a higher polymer feed and, thus, a thicker fiber. Based on the DoE data, regression analysis additionally enables a quantification of the effects of flow rate and drum speed on the fiber diameter (Figure S5d, Supporting Information).

**Figure 4:**
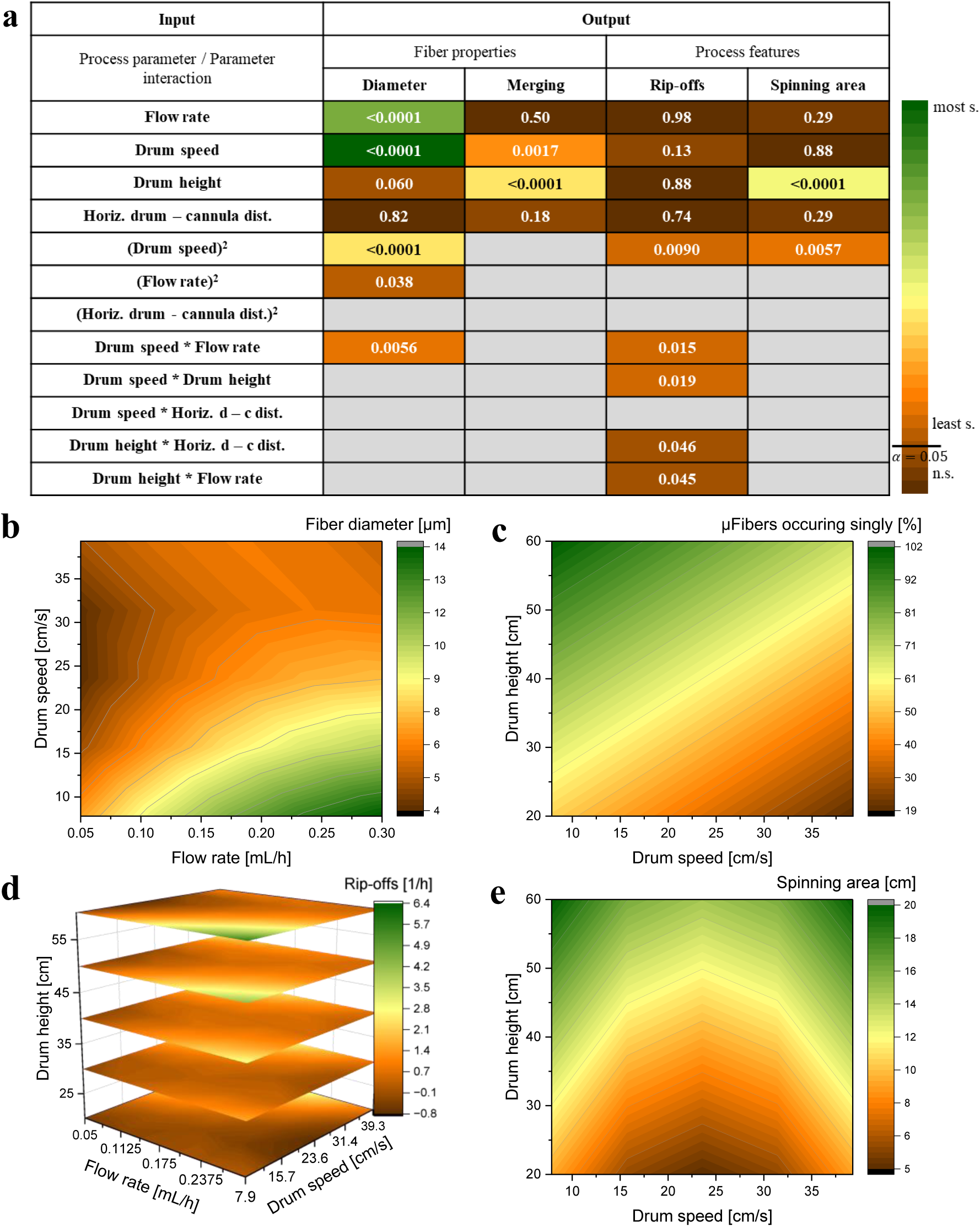
DoE I results. (a) Summary of the regression analysis, indicating the degree of influence of the tested process parameters or parameter interactions on respective output parameters (green: most, brown: least / not significant (s.)). A significance level of *α* = 0.05 was chosen for the analysis; respective *p*-values are given as numbers in the table. Interaction effects were stepwise removed from the model if not significant and are not listed unless they are significant for at least one of the output parameters (in this case, marked grey for output parameters for which they were not significant and removed from model). (b – e) Response surface plots modelled based on prediction equations obtained with regression analysis (equations see Figure S5 – S8, Supporting Information), yielding predictions for output parameters for untested input factor combinations within the defined parameter space. Only parameters with significant main or interaction effects are plotted on axes, non-significant parameters were set to level 0 values (compare coding in Figure 3b) in the modelling equation. Modelled output parameters: (b) fiber diameter, (c) amount of µfibers occurring individually, not merged, (d) average amount of fiber rip-offs per hour, (e) spinning area of fibers on the drum after spinning.

#### µFiber Merging

Regarding µfiber merging, the amount of µfibers occurring individually, not merged with others, experimentally varied between 90% (DoE run 1) and 20% (DoE run 27) (Figure S3c, Supporting information). A positive linear effect of drum height and a slightly weaker negative linear effect of drum speed were found to be main predictors for the amount of µfibers occurring as single fibers in a µfiber dispersion. This can be explained by a longer drying time of the filament before coming into contact with other fiber strands already collected when the drum is positioned higher above the cannula or pulls the filament more slowly. Consequently, minimal merging can be obtained at high drum heights and low drum speeds (Figure 4a and c; and Figure S6, Supporting Information). The flow rate, potentially influencing the efficacy of solvent evaporation from the filament through its influence on fiber thickness, did not exhibit a significant effect within the investigated parameter space (Figure 4a). Presumably, the filament core has less time to dry at higher flow rates while the filament surface dries more independently of the flow rate so that no influence on µfiber merging is observed. The polynomial equation fitted to the experimental data during regression analysis yields an *R*^2^ of 0.63. Any air movement around the spinning setup (positioned in a fume hood because of evaporating solvents) caused by fluctuations of laboratory air flow or movements of persons near the setup was identified as a noise factor not considered in the fits. Through air movement, the filament was visually observed to move laterally on the drum during spinning; this could influence fiber merging as the chance of fibers landing on top of each other before being completely dried is decreased with increasing lateral fiber movement.

As the efficiency of filament coagulation and drying, besides the investigated input parameters, also relies on the kinetics of the non-solvent-induced phase separation^[50]^ between PCL and ethanol, a different non-solvent for PCL could also influence the amount of µfiber merging. In this study, only ethanol has been used as non-solvent, but other non-solvents, e.g. dichloromethane, could be tested in future studies.

#### Fiber Rip-off Rate

During spinning, the fiber ripped off the drum occasionally, requiring manual collection of the extruded filament from the coagulation bath and its re-attachment to the drum to continue the process. To investigate how often the fiber, on average, ripped off per hour, and if an influence of the process parameters on this rip-off rate could be detected, all time periods of continuous spinning were logged and the fiber rip-off rate was calculated based on the observed fiber rip-offs and the total spinning time of the respective DoE run. Each run was continued throughout a full working day (see Figure S4, Supporting Information, for rip-off rates and total spinning times of each run).

Experimentally, the rip-off rate varied between 0 h^−1^ (DoE run 14 and 27), i.e. no rip-offs during the whole experiment, and ∼3 h^−1^(DoE run 2) (Figure S3d, Supporting information). None of the process parameters exerted a linear effect on the rip-off rate. However, a quadratic effect of the drum speed and slightly significant interaction effects between drum speed and flow rate, drum speed and drum height, flow rate and drum height, as well as drum height and horizontal drum – cannula distance (Figure 4a and d) were found during regression analysis. Due to the quadratic effect of the drum speed, desirable low rip-off rates can be achieved at middle drum speeds, e.g. 17 − 25 cm/s when all other process parameters are set to medium levels (Figure S7b, Supporting Information), while higher rip-off rates at lower or higher drum speeds indicate that the spinning process is less stable in these drum speed ranges. The interaction effect between drum speed and flow rate shifts the regime at which low rip-off rates are obtainable to lower drum speeds if low flow rates are employed, or to higher drum speeds if higher flow rates are used. A similar interaction effect is observed between drum speed and drum height (Figure S7e, left column of interaction plots, Supporting Information). Fitting of the experimental data during regression analysis yielded an *R*^2^ of 0.64. Further, it is observed that, with the obtained regression model, slightly negative rip-off rates are predicted at the boundaries of the investigated parameter space (Figure 4d). These observations indicate that noise factors, not considered in our model, influence how long a fiber can be spun continuously before ripping off. This aligns with our experimental observation that air movement around the spinning setup, as already described in the previous section, impacts the process stability in addition to the considered process parameters. Air movement can enhance the drying of the filament and thereby, on the one hand, increase its mechanical stability but, on the other hand, also make it more brittle and tenser,^[51]^ which could cause additional rip-offs. Shielding the setup with fabric or acrylic glass panels can help reduce air movement around the setup as a noise factor (see below in model validation).

#### Spinning Area of Fibers on the Drum

The spinning area, represented by the width of the drum section the fibers are distributed on after spinning, varied experimentally from 5.5 cm (DoE run 11, 27) to 17.0 cm (DoE run 2) (Figure S3e, Supporting information). It was found to be governed by a positive linear effect of the drum height, i.e. an increasing drum height increases the spinning area, and a quadratic contribution of the drum speed (Figure 4a and e; and Figure S8, Supporting Information). Similar to µfiber merging and fiber rip-off rate, air movement around the setup was visually observed to be a noise factor for the spinning area of the fibers on the drum (*R*^2^ = 0.67 of obtained fit). Additionally, the mechanical stability of the rotation axis of the drum constitutes a small noise factor as even slight movements of the rotation axis would broaden the spinning area on the drum. In general, a small spinning area of the fibers on the drum is desirable for fiber alignment, upscaling and parallelization of the spinning process.

#### Model Validation

To validate the models obtained by regression analysis, six validation experiments of fiber spinning were performed (four parameter combinations already tested within the DoE and chosen with utilized DoE software (JMP Pro 17.0.0, JMP Statistical Discovery LLC) as repetitions, two parameter combinations that were not part of the DoE and chosen by experimenters). The obtained fiber diameters matched the predicted ones in all validation runs (Figure S9a, Supporting Information) and µfiber merging was accurately predicted in two of the software-chosen repetition runs (runs 1 and 4 in Figure S9b, Supporting Information) and both experimenter-chosen runs (runs 5 and 6 in Figure S9b, Supporting Information), overall confirming the validity of the established models. During the validation runs, a shielding of the spinning setup with fabric was introduced to reduce the uncontrolled influence of air movement around the setup, resulting in desired lower fiber rip-off rates and smaller fiber distributions widths on the drum than predicted for all runs, except the first run in case of fiber rip-offs. Regarding the spinning area, the trend of the predicted areas is mostly mirrored in the experimental values, indicating that the effects of the process parameters established in the obtained model are still valid (Figure S9d, Supporting Information), while for the rip-off rates, the reduction of the air movement appears to be the most influential factor in the validation runs (Figure S9c, Supporting Information). In addition, it has to be noted that changes of ambient parameters such as temperature and humidity could also influence µfiber merging and the amount of fiber rip-offs due to changes of the fiber drying kinetics. These parameters were monitored during the conducted spinning experiments but not controlled in our setup, hence their influence is not considered in the model, but, as noise factors, they could cause deviations from the prediction model.

Overall, the conducted DoE approach revealed significant influences of the process parameters flow rate, drum speed and drum height as well as related quadratic and non-linear interaction effects on the investigated output parameters. In contrast, the positioning of the drum in front of, under or behind the drum (i.e. the horizontal drum – cannula distance) did not exhibit a significant main effect on the output within the investigated parameter space; only within non-linear parameter interactions with other parameters, slightly significant effects of this parameter were observed. As an increasing drum height both decreased undesired µfiber merging but increased (for upscaling) undesired broadening of the spinning area on the drum, a setting has to be chosen for this parameter depending on the focus of the output. Similarly, adjusting the drum speed is the strongest mean to set a desired fiber diameter but can simultaneously compromise µfiber merging, fiber rip-offs and the spinning area on the drum. With the correlative models presented here, the outcome for the investigated output parameters fiber diameter, µfiber merging and spinning area can be predicted when selecting specific combinations of process parameters; only the prediction efficiency of the rip-off rate model, as shown in the validation runs, is limited due to environmental noise factors impairing the DoE data.

### 2.3 Boundary Conditions for Spinnability and Upscaling

To test the limits of our spinning process regarding fiber production at extreme flow rates and drum speeds (i.e. spinnability) and the upscaling potential of the process, a second spinning setup (Setup II) was built, enabling higher drum speeds of 74 to 1048 cm/s (drum diameter: 5 cm, possible rotational drum speeds: 285 − 4000 rpm) compared to Setup I (2 − 42 cm/s). Furthermore, the reduction of air movement noticed in our Setup I as a noise factor was addressed by installing a ventilated shielding box around the spinning device (Figure S10, Supporting Information); the box is designed to fit on a standard laboratory bench below a suction arm for ventilation. The drum height and horizontal drum – cannula distance were fixed to 40 cm and −7 cm, respectively, in this setup (see Figure 3d).

First, the flow rate operability range for the spinning process was tested at the lowest possible drum speed of 74.6 cm/s by systematic variation of the flow rates between 0.01 and 0.40 mL/h. At flow rates below 0.05 mL/h, the extruded filament was too fragile to be collected at the cannula tip and placed on the drum. Attaching the fiber with a flow rate of 0.05 mL/h to the drum and subsequently reducing it to 0.01 mL/h was possible; however, the spinning was repeatedly disrupted by the filament breaking 1 − 2 min after the flow rate was decreased. Regarding the upper limit, no filaments could be attached to the drum at flow rates above 0.35 mL/h as the coagulation of the spinning solution in the coagulation bath was too slow to allow for the collection of a stable filament with tweezers. At 0.35 mL/h, attachment of the filament was feasible but, after a short period of spinning, excess spinning solution started flowing down the cannula tip while only a part of the solution, coagulated into a filament, was collected on the drum. This indicates that coagulation of the complete extruded spinning solution was not achieved at this flow rate. Similarly, the upper drum speed limit, tested at a stable flow rate of 0.30 mL/h, was found to be 523.6 cm/s (2000 rpm); at higher drum speeds, the filament would break after less than 2 min of spinning. Additionally, it was observed that, at the lowest flow rates or highest drum speeds tested, the air current created by the rotation of the drum close to its surface destabilized the filament, thereby further impairing successful attachment of the filament to the drum and, if initially attached, not supporting continuous spinning.

Based on these initial tests regarding spinnability, the effects of extreme spinning conditions on the fiber properties and process features were analyzed in a second DoE study (DoE II). Similar to DoE I, the output parameters fiber diameter, µfiber merging, fiber rip-offs and spinning area of fibers on the drum were screened. A full factorial design with additional center point (at least two replicates per point) was utilized, and flow rate (0.05 − 0.30 mL/h, matching the range employed in DoE I) and drum speed (300 − 2000 rpm, resulting in surface speeds of 78.5 − 523.6 cm/s, lower limit determined by limits of setup) were employed as input factors (Figure 5a). As the parameter combination of low flow rate (0.05 mL/h) and high drum speed (523.6 cm/s) did not allow a stable spinning process, the highest drum speed feasible for this flow rate was determined (144.0 cm/s) and the resulting parameter combination incorporated in DoE II, thereby also yielding the spinnability window marked in Figure 5a. Due to this adjustment, the DoE II design was not symmetric anymore, restricting the power of the DoE approach and precise predictability of the output parameters throughout the parameter space. Nonetheless, significant results regarding spinning at the boundary of the spinnability window and upscaling of µfiber production were obtained. Experimental data (Figure S11), detailed accounts on the polynomial regression analysis, effect summary plots, prediction profilers and resulting models can be found in the Supporting Information (Figure S12 – S17). As above, results for each output parameter, including comparisons to DoE I results, and the upscaling potential of the spinning process observed within DoE II are discussed in the following sections.

**Figure 5:**
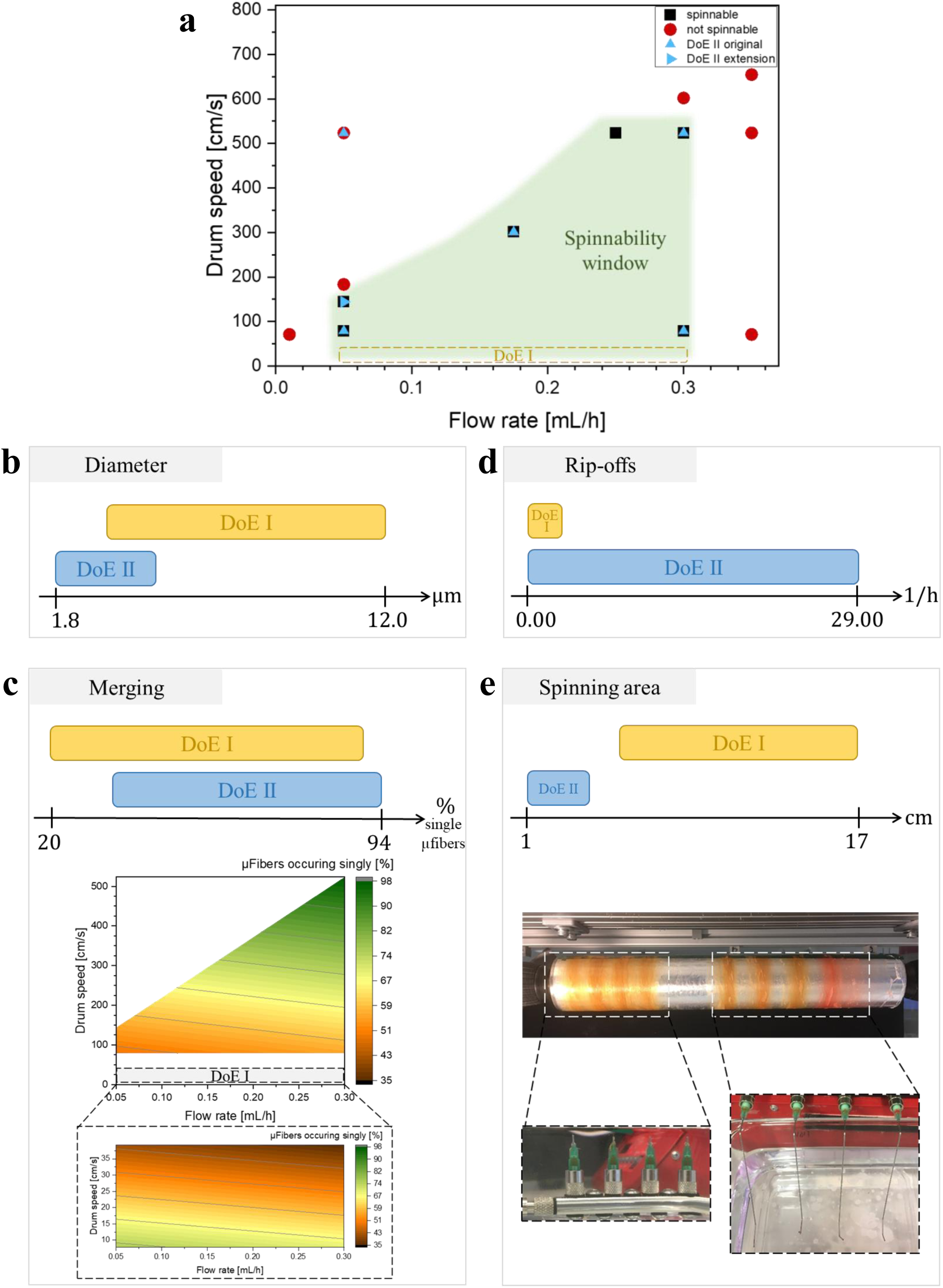
DoE II design approach and results. (a) Parameter combinations of flow rate and drum speed tested in initial spinnability tests and in DoE II (blue triangles). Parameter combinations allowing a continuous spinning process (> 2 min) (black squares) or prohibiting continuous spinning (red dots) establish the spinnability window (green). Parameter combination area investigated in DoE I marked with yellow dashed line. (b – e) Experimentally obtained output parameter ranges of DoE II in comparison to DoE I for (b) fiber diameter, (c) µfiber merging, additionally showing a comparison with DoE I, highlighting a trend change towards less µfiber merging at very high drum speeds used in DoE II; (d) average amount of fiber rip-offs per hour, (e) spinning area of fibers on the drum after spinning, additionally showing 1 − 2 cm broad spinning areas allowing parallelized fiber spinning with a multi-nozzle spinneret (bottom left) or several parallel syringes equipped with customized cannulas (bottom right).

#### Fiber Diameter

In DoE II, fiber diameters as small as 1.8 ± 0.4 µm (DoE run 12) (Figure S11b, Supporting Information) were obtained, extending the range of producible fiber diameters established in DoE I towards smaller ones (Figure 5b). As before, both flow rate and drum speed constituted significant effects on the fiber diameter (Figure S13, Supporting Information). The drum speed had a more pronounced effect on the fiber diameter at generally lower drum speeds used in DoE I compared to higher ones used DoE II (for comparison see Figure 4b: at a flow rate of 0.3 mL/h, decrease of fiber diameter from 14 to 6 µm by increasing drum speed from 8.4 to 37.7 cm/s (DoE I), versus Figure S17a, Supporting Information: at a flow rate of 0.3 mL/h, decrease of fiber diameter from 5 to 3 µm by increasing drum speed from 78.5 to 523.6 cm/s (DoE II)). This result coincides with the quadratic drum speed interaction observed in DoE I, which dampened the fiber diameter curve towards higher drum speeds (Figure S5b, Supporting Information). Due to the very high drum speeds used in DoE II, the thickest fiber diameter achievable with Setup II / in DoE II was 4.9 ± 0.6 µm (DoE run 13), compared to 12.0 ± 2.1 µm in DoE I.

#### µFiber Merging

Regarding µfiber merging, a significant trend change was noticed: while at generally lower drum speeds employed in DoE I, the amount of single, non-merged µfibers decreased with increasing drum speed, this trend was inversed in the range of 40 to 80 cm/s, yielding more individual µfibers at extremely high drum speeds in DoE II (Figure 5c; and Figure S14, Supporting Information). This can be explained by an interplay of multiple effects at very high drum speeds: First, an increased drum speed yields more strongly stretched fibers with a smaller fiber diameter, facilitating the complete solvent evaporation from the filament before being collected on the drum. Second, through a faster axial movement of the filament towards the drum at higher drum speeds, an increased air movement near the filament surface facilitates the removal of solvent molecules evaporated from the fiber surface, thereby enhancing the solvent evaporation. This effect might be further reinforced by a strong air flow near the drum surface caused by the drum rotation. In contrast to this, in DoE I, higher drum speeds led to increased fiber merging as the effect of shorter drying times seemed to dominate within the parameter window tested there.

#### Fiber Rip-off Rate

To observe how often the fiber rips off during the spinning process, spinning of each DoE run was continued for 2 to 5 h (half a working day), and the average amount of fiber rip-offs per hour was calculated similar to DoE I (Figure S11d and Figure S12, Supporting Information). In DoE II, continuous spinning without the fiber ripping off (i.e. fiber rip-off rate of 0 h^−1^) during the whole experiment was achieved in three different runs (DoE runs 10, 13, 14). In all of these runs, the lowest drum speed of 79 cm/s was employed, while the highest flow rate of 0.3 mL/h was applied in two of those runs and the lowest one of 0.05 mL/h in the third one, indicating a stable process throughout the whole flow rate range at low drum speeds. In contrast to this, higher rip-off rates of up to 29 h^−1^ (DoE run 2, flow rate: 0.175 mL/h, drum speed: 301 cm/s, Figure S11d, Supporting Information) were obtained at higher drum speeds in DoE II. Therefore, despite the reduction of the ambient air movement, which was assumed to be a noise factor for the rip-off rate in DoE I, by implementing the shielding box around Setup II, the range of rip-off rates observed in DoE II is ten times broader than in DoE I (Figure 5d). This indicates a decreasing process stability at more extreme drum speeds (Figure S15 and S17c, Supporting Information). Besides extreme fiber stretching, the air current created by the rotation of the drum, assumedly promoted more frequent fiber rip-offs at very high drum speeds.

#### Spinning Area of Fibers on the Drum

In DoE II, spinning areas on the drum varied between 1 and 4 cm (DoE runs 11, 12 and 13, Figure S11e, Supporting Information). Overall, significantly smaller spinning areas were obtained compared to DoE I (Figure 5e). This is mainly attributed to the reduced air movement through shielding in Setup II, as uncontrolled environmental air movement was visually observed to be an influential noise factor broadening the spinning area in Setup I through lateral fiber movement on the drum. Smaller spinning areas are desired for upscaling as they enable parallelized spinning of multiple filaments as demonstrated in the bottom images of Figure 5e. Here, eight filaments were fabricated in parallel, one half with a multi-nozzle spinneret connected to a single syringe containing spinning solution, the other half with four separate customized cannulas, each connected to a separate syringe. The separate cannulas allowed for distinct spacing of the fibers on the drum, which is superior if the fibers produced from each cannula should be collected and post-processed separately. In contrast, the multi-nozzle spinneret enabled closer spacing and a facilitated preparation of only one syringe. However, the spinning areas of the fibers extruded through the different nozzles of the multi-nozzle spinneret slightly overlap, allowing only a collective post-processing of all four filaments. Further, a slightly higher risk of the spinning being disrupted by entanglement of two different filaments, causing them to tear off, was observed when spinning with the multi-nozzle spinneret.

#### Upscaling of µFiber Production

Regarding upscaling, fabrication of µfibers with Setup II, compared to initial fabrication with Setup I, permits an increase in the production rate due to more efficient parallelization with smaller spinning areas and higher possible drum speeds. Exemplary, for the production of µfibers with a diameter and length of 5 µm and 50 µm, respectively, the production rate could be increased more than three times from 140 ⋅ 10^6^ with Setup I (spinning fibers from 3 needles in parallel) to 470 ⋅ 10^6^ fibers/work day with Setup II (8 fibers in parallel). Notably, upscaling of the production of µfibers thicker than 5 µm is restricted with the current Setup II as the employed motor does not permit drum speeds below 285 rpm (with current drum diameter resulting in 74.6 cm/s surface speed) needed for the fabrication of thicker fibers. If thicker µfibers are found to be advantageous for specific 3D cell culture models, upscaling here can be achieved in the future by implementing either a gear or different motor or a drum with smaller diameter in Setup II to enable lower surface drum speeds in a shielded environment.

### 2.4 Magneto-Responsiveness and Fluorescence of µFibers

To render the µfibers magneto-responsive and, thus, alignable in an external magnetic field, SPIONs, coated with fatty acid for enhanced dispersibility in chloroform (shell: 23 ± 1 wt% of the particles, based on thermogravimetric analysis), were included in the spinning solution. Two different preparation strategies for the spinning solution, containing PCL, chloroform, a fluorescent dye and SPIONs, were compared: in the first approach, PCL was dissolved in chloroform containing fluorescent dye under continuous stirring, yielding a viscous solution. SPION stock dispersion (in chloroform) was added and the solution was mixed using an ultrasonic homogenizer with horn tip as previously reported.^[23]^ In resulting spinning solutions, aggregates of the black SPIONs were visible after ultrasonication, sedimenting when filled into a syringe (Figure S18a, Supporting Information). Prolonging the sonication time did not yield a disruption of the aggregates; additionally, extensive heating of the solution despite cooling of the solution vial in an ice bath during sonication was noticed, causing an uncontrolled evaporation of chloroform. µFibers prepared with this solution exhibited SPION aggregates in the range of several micrometers near the fiber surface, imaged by energy dispersive X-ray spectroscopy analysis (Figure S18b, Supporting Information). In a second spinning solution preparation approach, chloroform, fluorescent dye and SPION stock dispersion were mixed by pipetting and vortexing. Then, PCL pellets were added, the solution was vortexed for 1 min and treated on a roller mixer overnight to ensure full dissolution of the polymer. Resulting spinning solutions appeared homogenous and corresponding µfibers exhibited a homogenous distribution of iron content (Figure S18c and d, Supporting Information). Consequently, the second approach was used for spinning solution preparation throughout this study to ensure a homogeneous SPION distribution in the fibers for optimal magneto-responsiveness in an external magnetic field (Figure 6a).

**Figure 6:**
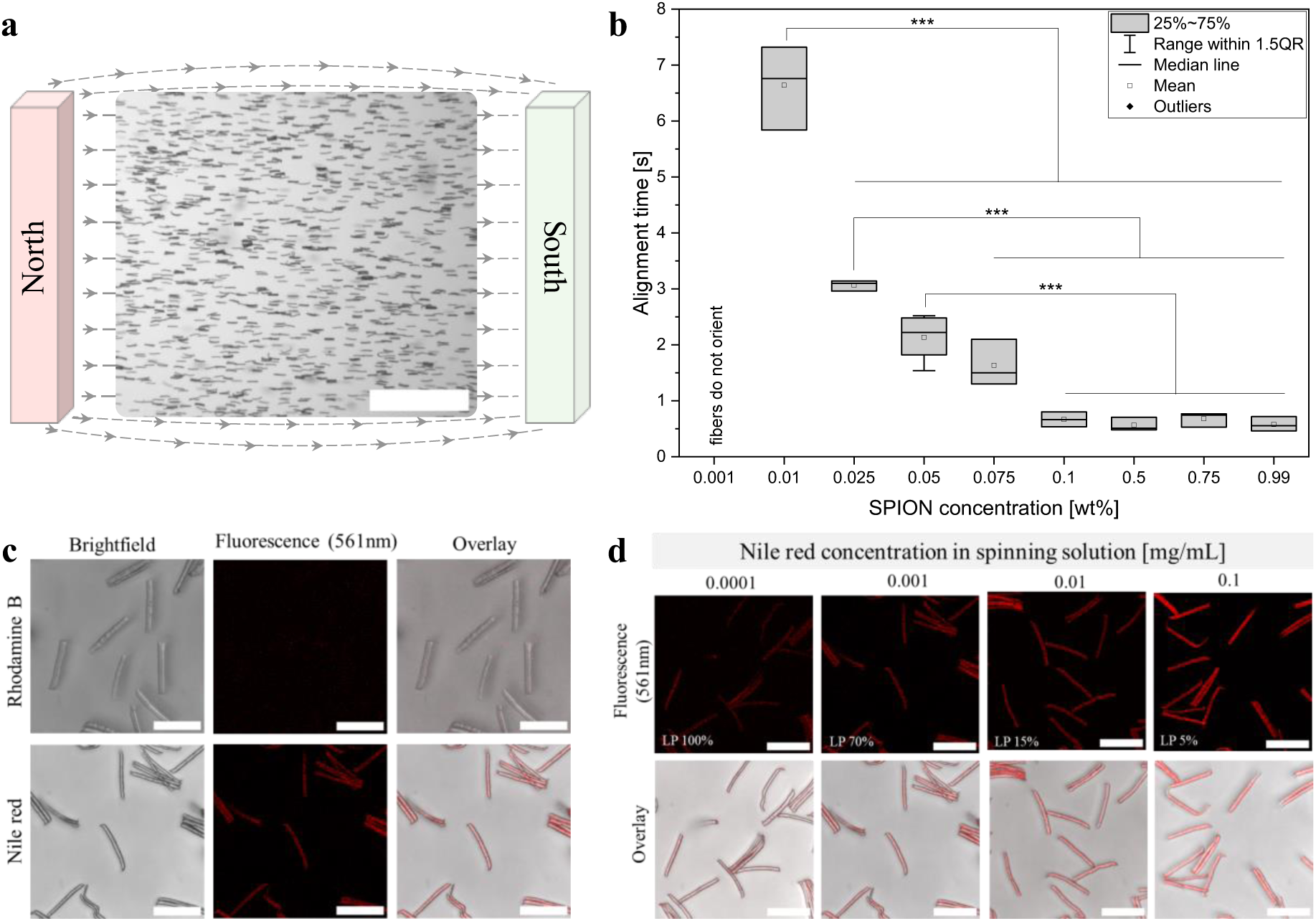
(a) µFibers dispersed in water, aligned by an external magnetic field. Scale bar: 500 µm. (b) Alignment time needed to align µfibers dispersed in water in an external magnetic field (40 mT) depending on their SPION concentration (given as wt% in relation to the polymer weight); \*\*\**p* < 0.001 (ANOVA, post-hoc Bonferroni test). (c) Fluorescence confocal microscopy images of µfibers with RhB (0.1 mg/mL in spinning solution) or nile red (0.001 mg/mL). The laser power during imaging was set to 70%; other imaging settings were kept constant. Scale bars: 50 µm. (d) Fluorescence microscopy images of µfibers spun with concentrations of 0.0001 – 0.1 mg/mL nile red in spinning solution. The laser power (LP) needed to acquire the images varied between 100% and 5%, as noted at the bottom of the fluorescent images; other imaging settings were kept constant. Scale bars: 50 µm.

To assess the influence of the SPION concentration incorporated in the µfibers on their ability to align within a weak magnetic field (40 mT), the alignment time of µfibers fabricated with different SPION concentrations was measured in water (Figure 6b). The time needed for alignment is relevant for intended applications of µfibers as anisotropic building blocks in 3D hydrogel cultures as it has to be shorter than the hydrogel gelation time. SPION concentrations of 0.99 wt% down to 0.1 wt% yielded an quasi-instantaneous alignment of the µfibers within 1 s; further reduction of the concentration resulted in increased alignment times of up to 6.7 ± 0.8 s at 0.01 wt%. At very low SPION concentrations of 0.001 wt%, no alignment was observed. These results indicate that the SPION content can be reduced down to 0.1 wt% or 0.01 wt%, depending on the desired alignment time, employed magnetic field, and the properties of the surrounding matrix chosen for embedding the µfibers to form a 3D Anisogel. In the case of 1 vol% of µfibers inside the Anisogel, both concentrations would be significantly below known toxicity levels (∼1.5 mM of maghemite SPIONs in direct cell contact^[52]^) and thus be biocompatible.

Incorporation of rhodamine B (RhB) or nile red in the spinning solution was investigated to render the µfibers fluorescent and, thus, enable facile analysis of µfiber alignment and distribution inside 3D Anisogels by confocal fluorescence microscopy. RhB was successfully used as a stain without covalent binding for different polymeric fibers before;^[23,53]^ however, no sufficient µfiber fluorescence could be achieved with RhB concentrations of up to 0.1 mg/mL in the spinning solution (Figure 6c, top; and Figure S19a, Supporting Information) in this study. Due to its solubility in ethanol,^[54]^ RhB is partly lost due to diffusion from the extruded filament into the coagulation bath during the spinning process. Although the retention time of the filament in the coagulation bath is very short, a red-pink coloration of the coagulation bath after spinning indicated RhB diffusion into the bath. Additionally, it is hypothesized that the RhB fluorescence is partly quenched by interaction with SPIONs in the spinning solution.^[55]^ In contrast, nile red, exhibiting a strong hydrophobicity^[56]^ and being recently applied as a detection dye for hydrophobic microplastics,^[57]^ was successfully integrated into our fibers based on physical interactions and yielded good fluorescence after fiber cutting and µfiber purification (Figure 6c, bottom; and Figure S19b, Supporting Information). After testing of different nile red concentrations in the spinning solution (0.0001 − 0.1 mg/mL), a concentration of 0.01 mg/mL, yielding sufficient fluorescence at moderate laser power, was used as a standard dye concentration for µfiber production (Figure 6d).

Possible cell toxicity of µfibers with the highest used SPION concentration (∼1 wt%) and optimized nile red dye concentration (0.01 mg/mL in the spinning solution) was tested in a cell viability assay. µFibers did not decrease the cell viability and were found to be biocompatible (Figure S20, Supporting Information).

### 2.5 µFibers as Guiding Elements in a 3D High-Throughput Vasculogenesis Model

To investigate the influence of our µfibers on cell alignment in a HT compatible 3D cell culture model, µfibers (5.6 ± 1.2 µm diameter, 51.1 ± 3.3 µm long) were implemented in a 3D vasculogenesis model produced and maintained with an automated fluid handling and cell culture system (Explorer^™^ G3 workstation, including JANUS^®^ G3 and Opera Phenix™ Plus high-content imaging system, Revvity), established by Dennison, Fusenig and Groennert *et al*.^[58]^ Here, a co-culture of HUVECs and MSCs was embedded in growth factor-affine multi-armed poly(ethylene glycol) and sulfonated glycosaminoglycan (heparin) based hydrogels (starPEG-sGAG hydrogels) displaying RGD peptides for cell adhesion and matrix metalloprotease (MMP)-cleavable peptide crosslinks allowing for cellular remodeling.^[59]^ The biorthogonal Michael type click reaction utilizing cysteine-containing peptides and maleimide-pre-functionalized heparin was optimized for the automated gelation in the low-volume 384 well plate format. Cell densities, cell ratios of co-cultured cells, the concentration of co-embedded growth factors, and cell culture medium composition were systematically optimized for longest HUVEC sprout formation or most efficient (i.e. lower growth factor concentrations, more cost-effective) cultivation conditions.^[58]^

For implementation of our µfibers into this established vasculogenesis model, a device creating a weak quasi-linear magnetic field throughout a HT format well plate, and being compatible with deck layouts of automated fluid handling systems, constituted a key requirement to enable the embedding of µfibers in an aligned manner. A custom-made magnetic well plate lid suitable for a 96 well plate was developed (Figure S21a, Supporting Information); it can be mounted onto a well plate while the well plate itself is placed in a conventional well plate holder of a liquid handling workstation (here: JANUS^®^ G3, Revvity). To match the magnetic lid format and simultaneously retain a low volume culture model, the model established by Dennison, Fusenig and Groennert *et al.* was adapted to a specialized 96 µwell plate, in which a hydrogel can be casted in an inner well with a volume of 10 µL (Figure 7a), while the magnetic lid, with magnets positioned on neighboring wells, created a quasi-uniform magnetic field in the culture well (every second row of the well plate usable for culture). Pipetting protocols were further adjusted to the incorporation of the µfiber dispersion inside the starPEG solution as precursor solution for the hydrogels; cells and growth factor supplementation were kept in the sGAG stock solution, and mixed with the starPEG stock solution in a 96 well preparation plate. After automated transfer of the mixed hydrogel precursor solution to the final 96 µwell plate and a latency time (10 min) to allow for complete hydrogel gelation, the magnetic lid was replaced with a standard well plate lid. Medium changes, brightfield and confocal fluorescent imaging during culture were performed in an automated manner with a robotic workstation (Explorer^™^ G3, including Opera Phenix™ Plus, Revvity).

**Figure 7:**
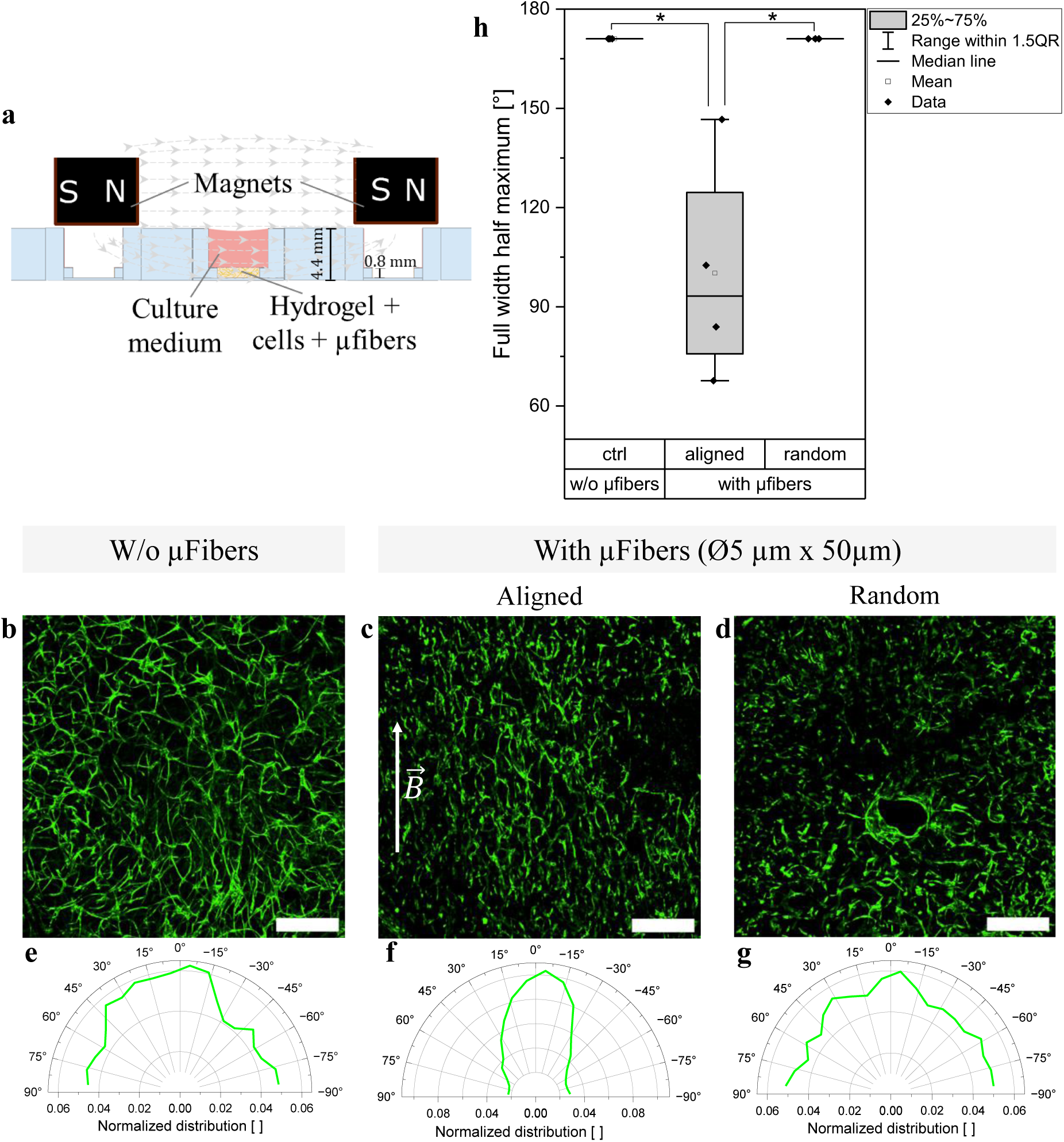
(a) Setup for cell culture model fabrication. A starPEG-sGAG hydrogel, pre-loaded with growth factors and RGD-adhesive peptides, containing HUVECs, MSCs and µfibers is casted into a 96 µwell plate equipped with a custom-made magnetic well plate lid (magnetic field indicated by light grey arrows). The magnetic lid is replaced with a common well plate lid after hydrogel gelation. (b – d) Exemplary confocal fluorescence microscopy images of HUVECs (GFP expressing, green) cultured with MSCs (not stained, ratio 95:5) and (b) without, (c) with magnetically aligned or (d) with randomly oriented µfibers in a starPEG-sGAG hydrogel at DIV13 (maximum projection of 106 µm thick z-stack within hydrogel). Scale bars: 500 µm. (e – g) Corresponding angular orientation of HUVEC structures. (h) Full widths at half maximum (FWHMs) obtained from angular orientation of HUVEC structures using maximum projections of z stacks; \**p* < 0.05 (ANOVA, post-hoc Bonferroni test).

Vascular sprouts grown in hydrogels with µfibers exhibited a preferential orientation coinciding with the orientation of the magnetically aligned µfibers (Figure 7c; orientation of µfibers shown in Figure S21b – d, Supporting information). In comparison, isotropic growth was observed in hydrogels without µfibers (Figure 7b) or hydrogels with randomly oriented µfibers (no magnetic lid used during hydrogel formation) (Figure 7d). The corresponding full width at half maximum (FWHM) of the aligned HUVEC structures (100 ± 34 °), yielded by image analysis of maximum projections created from z stacks, is significantly smaller than the FWHM of the isotropic structures (171 ± 0 °, highest possible FWHM here due to radial increments chosen in image analysis as described in Experimental Section, Supporting Information) (Figure 7e – h). Additionally, randomly orientated µfibers seemed to inhibit the vascular sprout formation (Figure 7d).

These results clearly indicate that the obtained µfibers are a powerful tool to influence the growth direction of endothelial cell sprouting in vasculogenesis models, and are suitable for automated HT tissue culture. In future studies, the effects of varying µfiber dimensions and concentrations on the vascular network formation, in particular filament lengths, branching points, the time frame necessary to form aligned structures, as well as the integration of the vascular networks in perfusable platforms^[60]^ will be investigated. Further, the implementation of our µfibers allows to prospectively compare responses of aligned vasculogenesis models in dose-dependent vasculogenesis inhibitor drug testing to already established isotropic models,^[58]^ paving the way towards improved HT tissue models for *in vitro* drug testing, improving personalized medicine and reducing animal testing.

## 3 Conclusion

Short fibers are versatile building blocks to introduce anisotropy in 3D hydrogel cell culture models, bioinks or injectable platforms. Therefore, a controlled fabrication of short fibers with precisely tailored dimensions and in quantities suitable for high-throughput applications are required.

In this study, we introduced a scalable semi-continuous production process, comprised of wet-dry fiber spinning and subsequent cryosectioning, to fabricate short magneto-responsive fluorescent PCL µfibers with diameters in the low micrometer range and variable lengths with narrow size distributions. To enable a precise production of µfibers with specific dimensions in the future, correlative prediction models for fiber properties and process features were established by employing DoE approaches for experimental planning and analysis. With upscaling of the process, µfiber production rates of up to nearly half a billion µfibers per work day were achieved, and boundary parameter conditions of the spinning process were investigated. Magnetically aligned µfibers implemented into a previously established, automated high-throughput vasculogenesis model triggered oriented, anisotropic vascular sprout formation, showcasing the potential of our µfibers as building blocks for anisotropic tissue models and high-throughput applications. Overall, our established µfiber fabrication process offers a versatile, scalable platform for the production of short PCL µfibers, and, prospectively, can be adapted for fabrication of other polymeric fibers to be used in novel tissue engineering solutions aiming at mimicking naturally anisotropic human tissues.

## 4 Experimental Section

Experimental details can be found in the Supporting Information.

## Supporting Information

Supporting Information is available online or from the author.

## Acknowledgements

This work was supported through funding of the Senatsausschuss Wettbewerb (SAW) Leibniz-Transfer project *µ*TISSUE*fab* (A.A.M., R.N., M.F., N.R.D., C.W., L.D.L.) and Leibniz Health Technologies Short Term Scientific Mission (STSM) funding (A.A.M., R.N.). Further support of the Deutsche Forschungsgemeinschaft (DFG GRK2415 ME3T), the Government of Canada’s New Frontiers in Research Fund (NFRFT-2020-00238: Mend the Gap) and the European Research Council (ERC – Heartbeat, Grant Agreement No. 101043656) is acknowledged. The authors thankfully acknowledge the support of the RWTH Aachen Interdisciplinary Center for Clinical Research (IZKF) in cryosectioning experiments and the support of the RWTH Aachen AME workshop in building the spinning devices. Additionally, the authors thank Dr. Rostislav Vinokur for assistance in building the spinning device controllers and the rail for stepless drum height variation, Stefan Hauk for assistance with SEM and EDX measurements and Nina Schöling for assistance with fiber spinning experiments and µfiber purification.

## Data Availability Statement

Data that support the findings of this study are available in the Supplementary Information of this article. Automation scripts, JMP scripts or further data are available from the corresponding author upon reasonable request.

