## Supplementary Information for "Fabrication of Short Polymeric µFibers as Building Blocks for Anisotropic High-Throughput Compatible 3D Tissue Models"

A. A. Meyer, M. Harmeth, J. Graeve, K. Rengel, M. Keskin, T. Hülsmann, T. Haraszti, L. De Laporte
Institute of Technical and Macromolecular Chemistry (ITMC), Chair of Macromolecular Materials for Medicine, RWTH Aachen University, 52074 Aachen, Germany

M. Fusenig, N. R. Dennison, U. Freudenberg, C. Werner
Leibniz Institute of Polymer Research Dresden e.V., Max Bergmann Center of Biomaterials, 01069 Dresden, Germany

C. Werner
Center for Regenerative Therapies Dresden and Cluster of Excellence Physics of Life, Technische Universität Dresden, 01307 Dresden, Germany

L. De Laporte
Advanced Materials for Biomedicine (AMB), Institute of Applied Medical Engineering (AME), University Hospital RWTH Aachen, Center for Biohybrid Medical Systems (CMBS), 52074 Aachen, Germany

**Experimental Section**

*Materials:* For wet-dry spinning and subsequent µfiber production, chloroform ($CHCl_3$, 99.8%, Fisher Chemical), ethanol (EtOH, 99.8%, Fisher Chemical), ultrapure water ($H_2O$, MilliQ grade, inhouse filtration (Purelab Chorus I system, ELGA)), poly(ε-caprolactone) (PCL, 80 kDa, Sigma Aldrich), superparamagnetic iron oxide nanoparticles (SPIONs, EMG1200 (fatty acid shell), Ferrotec), rhodamine B (RhB, 97%, VWR), nile red (NR, Carl Roth), and cryosectioning medium Tissue-Tek O.C.T. Compound (OCT, Sakura, Weckert Labortechnik) were used. For cell culture experiments, Dulbecco's Modified Eagle medium (DMEM, Gibco), endothelial cell growth medium (ECGM, PromoCell) and, as medium supplements, fetal bovine serum (FBS, Capricorn Scientific), low-serum growth medium supplement mix (Growth Medium SupplementMix, PromoCell), antibiotics-antimycotic (AmB, Gibco) and penicillin/streptomycin (P/S, Sigma Aldrich) were used to culture L929 mouse fibroblasts (Deutsche Sammlung von Mikroorganismen und Zellenkulturen GmbH, DSMZ ACC-2), GFP-expressing human umbilical vein endothelial cells (HUVECs, Angioproteomie) and primary human mesenchymal stromal cells (MSCs). TrypLE Express Enzyme (1X, Gibco), fibronectin (20 µg/mL in PBS, prepared as published in Dennison, Fusenig, Groennert et al.[1]), LIVE/DEAD Cell Imaging Kit (488/570, Invitrogen) were used for trypsinization, live / dead staining and coating of tissue culture flasks, respectively. For hydrogel preparation, four-armed poly(ethylene glycol) (PEG) end-functionalized with a matrix metalloproteinase-responsive peptide (starPEG-peptide conjugates, 15.1 kDa, custom-synthesized[2] and provided by IPF Leibniz Institute of Polymer Research, Dresden), maleimide-functionalized heparin (sGAG, 15 kDa, six maleimide groups per heparin molecule, custom-synthesized[1] and provided by IPF Leibniz Institute of Polymer Research, Dresden), fibronectin-derived RGDSP peptide (990 Da, custom-synthesized[3] and provided by IPF Leibniz Institute of Polymer Research, Dresden), VEGF (Human VEGF 165 IS, Miltenyi Biotec), SDF1α (Human SDF-1α / CXCL12, Miltenyi Biotec) and FGF2 (Human FGF-basic / FGF-2 / bFGF, 154 aa, PeproTech) were used.

*Wet-dry Fiber Spinning*: For spinning solution preparation, based on a pre-weight amount of PCL, $CHCl_3$, fluorescent dye (RhB / NR) stock solution ($1\ mg/mL$ in $CHCl_3$) and SPION stock dispersion ($100\ mg/mL$ in $CHCl_3$, ultrasonicated for $1\ min$ before use) were pipetted into a glass vial and vortexed. The pre-weighed PCL was added, the solution vortexed for $60\ s$ and treated on a roller mixer ($30 - 50\ rpm$) overnight for complete dissolution of PCL. If not stated otherwise, spinning solutions contained $17\ wt/vol\%$ PCL, $0.01\ mg/mL$ NR and $1.70\ mg/mL$ SPIONs (equals $0.99\ wt\%$ of solid polymer content). The spinning solution was transferred into a glass syringe (TLL1002 / 1005, Hamilton) equipped with a custom-bent U-shaped, blunt cannula (Sterican $21\ G$, $120\ mm$, B.Braun) and placed in a syringe pump (AL-1000, World Precision Instruments). The pump was positioned behind an EtOH bath so that the U-shaped cannula was immersed in the bath, the cannula tip pointing upwards circa $2\ cm$ underneath the EtOH surface. For spinning, extrusion of the spinning solution was started. The extruded filament was picked up with a tweezer at the cannula tip, slowly drawn upwards and placed on a rotating aluminum drum (custom-built setup) situated above the EtOH bath, on which the filament was collected in an aligned manner. The drum was previously swathed in aluminum foil and coated with a dried layer of OCT. When the fiber collection was finished, the spun fibers were coated with a layer of OCT and left to dry overnight.

Spinning was conducted with two different rotating drum setups in this study. Setup I was placed inside a fume hood; the process parameters drum speed, drum height (measured from cannula tip) and horizontal drum – cannula distance could be varied in the ranges of $10$ to $200$ revolutions per minute (rpm), $5$ to $80\ cm$ and $-10$ to $+15\ cm$ (i.e. cannula in side view left or right of clockwise rotating drum, as depicted Figure 3d), respectively. The drum diameter was $4 \pm 0.1\ cm$, and the drum was $30\ cm$ long. Setup II was installed in a custom-built acrylic glass box with closable doors ($100\ cm$ wide, $81\ cm$ high, $60\ cm$ deep), ventilation was enabled through small holes at the edge of the ceiling via a suction arm above. The drum diameter was

$5 \pm 0.1$ cm, and the drum was 30 cm long. Here, drum speed could be varied from 285 to 4000 rpm, drum height and horizontal drum – cannula distance were fixed to 40 cm and $-7$ cm, respectively. For both setups, the flow rate as a further process parameter was controlled by the syringe pump and varied between 0.01 and 0.40 mL/h.

For a coherent comparison of the influence of the drum speed on fiber and process properties between experiments conducted with the two different setups, the linear speed at the drum surface $v$ (in cm/s) was calculated based the rotational speed $\omega$ (which was programmed with the drum controller for each experiment in rpm) and the respective drum diameter $d$ according to Equation (1). For all analyses, the surface drum speeds were used.

$$v = \frac{2\pi(d/2)\omega}{60} \tag{1}$$

*Cryosectioning and µFiber Purification*: Produced fibers were processed into short µfibers as reported previously.[4] Briefly, spun fibers were harvested from the drum by cutting the aluminum foil holding the fiber-OCT-film open along the drum. The film was peeled off the aluminum foil and sectioned into smaller pieces which were embedded into a silicon cryomold (3 cm x 5 cm) with OCT and frozen at $-20$°C; it was ensured that the pieces were stacked with fiber alignment along the long side of the cryomold. The frozen fiber-OCT-block was sectioned perpendicular to the fiber alignment into, if not stated otherwise, 50 µm thick slices using a cryostat microtome (Leica CM1950, equipped with FEATHER C35 microtome blades) at $-22$°C. Harvested sections were collected in a centrifuge tube, immersed in 50 mL water and treated on a roller mixer for at least 1 h at room temperature to dissolve the OCT. Following, the µfibers were purified by washing four times with water, with intermediate centrifugation (4500 rcf, 10 min) accelerating µfiber sedimentation.

For further analysis of fiber cross-sections, 1 or 5 µm thick sections were cut from respective fiber-OCT-blocks and collected either directly on glass slides (for fluorescence microscopy) or on glass slides covered with double-sided tape (for scanning electron microscopy).

*Design of Experiments (DoE) Experimental Planning and Data Analysis*: JMP Pro 17.0.0 (JMP Statistical Discovery LLC) software was used to design, model and evaluate two experimental series, DoE I and II. All considered factors were defined as continuous; for simplification of experimental planning and presentation, the real parameter values were also normalized to coded values representing the levels of the parameters chosen in specific DoE runs as typically done in DoE analyses (coded levels see Figure S3a and Figure S11a). For DoE I, a central composite design (star points with $\alpha_{\mathrm{CCD}} = 2$) with the four input factors flow rate, drum speed, drum height, horizontal drum – cannula distance and four replicates of the center point was chosen. Spinning of each run was conducted for a full working day. Polynomial regression analysis considering main, quadratic and non-linear interaction effects (of up to two input parameters, computing second order polynomials) was used to establish correlative prediction models for the four output parameters fiber diameter, µfiber merging, fiber rip-off rate and spinning area of fibers on the drum after spinning. To allow differentiation between significant and non-significant effects during regression analysis, a significance level of $\alpha = 0.05$, typically used in DoE analyses, was chosen. Effects with a probability value $p < \alpha$ were considered significant. After regression analysis, four additional replicates of runs with parameter combination settings already used in DoE runs (chosen with the JMP software) and two additional runs with factor settings that were not part of the original DoE were performed to evaluate the prediction model yielded by regression analysis. For DoE II, a full factorial design with additional center point was created for the two factors flow rate and drum speed, investigating boundary conditions of up to 13-fold higher drum speeds than used in DoE I. All runs were replicated once and conducted for $2 - 5$ h (half working day). Regression analysis was performed similar to DoE I.

Drum speed levels for both DoE approaches were determined with the JMP software in rpm as this unit was used to program the drum controllers; calculation of respective surface speed units for analysis was performed subsequently as described above (Equation (1)).

*Fiber Diameter, Fluorescence and Stacking*: Brightfield and fluorescent images of µfibers and µfiber cross-sections were recorded with a confocal fluorescence microscope (Leica TCS SP8) with 10x and 63x air objectives and a 561 nm laser excitation wavelength. The mean fiber diameter and length (given with respective standard deviations in main text) were determined by measuring the µfiber width and length in 63x magnification images using ImageJ (version 1.54f); at least 40 µfibers were measured per sample. The µfiber merging was quantified by counting µfibers with the ImageJ Cell Counter plugin. µFibers were counted in two categories: single µfibers, and µfibers which are merged and thus forming stacks of two or multiple fibers. At least 52 µfibers were counted per sample.

*Calculation of Fiber Rip-off Rate:* During fiber spinning, it was logged how often the fiber ripped off by itself and how long continuous spinning was possible in between. If a fiber was detached manually from the drum at the end of the experiment day, this event was not counted as a fiber rip-off, but the last continuous spinning period was clogged up to this time point. The fiber rip-off rate was calculated based on the amount of observed fiber rip-offs $x$ during the experiment day and the sum of continuous spinning times $t_i$ constituting the total spinning time of the experiment:

$$Rip - off\ rate = \frac{x}{\sum t_i} \quad (2)$$

*Fiber Morphology and SPION Distribution on Fiber Surface:* Fiber morphology was analyzed with field emission scanning electron microscopy (FE-SEM) (Hitachi S4800 with secondary electron (SE) detector). For imaging the fiber surface, samples of non-OCT-coated spun fibers were harvested from the collector after spinning; for cross-sections, respective slices collected on glass slides during cryosectioning were utilized. Samples were transferred onto SEM sample holders and coated with carbon (5 nm) using a sputter coater (Leica EM ACE600). SEM images were acquired at an acceleration voltage of 5 kV.

With energy dispersive X-ray spectroscopy (EDX) (Hitachi SU9000 with SE detector for SEM images and Oxford Instruments Xmax 80 detector with Aztec software for corresponding EDX images), the distribution of iron on the µfiber surface was mapped. Aqueous µfiber dispersions were pipetted onto copper grids with carbon films (AGS160-3H Carbon Films on 300 Mesh Grids Copper, Agar Scientific), dried overnight and sputter-coated with carbon (3 nm). SEM images were recorded at an acceleration voltage of 30 kV and respective EDX images with 15 kV. Per sample, five µfibers were analyzed.

*Determination of SPION Core to Shell Ratio via Thermogravimetric Analysis:* SPIONs utilized in this study exhibit an iron oxide core and an oleic acid shell. As the manufacturer did not give exact information about the core to shell ratio, thermogravimetric analysis (TGA) was conducted for analysis. At least 10 mg sample were weight out into an aluminum oxide crucible and placed in the thermogravimetric analyzer (NETZSCH TG 209 C, NETZSCH-Gerätebau GmbH, Germany). The organic components of the sample were decomposed under nitrogen atmosphere (oleic acid decomposes under nitrogen atmosphere without residues[5]) during a temperature ramp from 70 to 950°C with a heating rate of 10 °C/min; the final temperature was maintained for 30 min. The residual weight after the measurement was used to calculate the iron oxide content of the SPIONs.

*µFiber Orientation in External Magnetic Field*: Videos of the µfiber orientation in an external magnetic field were recorded with an inverted light microscope (Motic AE2000 and Flir camera FL3-U3-13Y3M-C with FlyCap2 2.13.3.31 Point Grey Research software) with a frame rate of 25, 50 or 150 frames per second (depending on the time scale of fiber orientation) using an 4x objective. 20 µL of aqueous µfiber dispersion were pipetted onto a glass slide placed in a custom-made magnetic slide holder (40 mT) while recording a 10 s to 2 min video (duration depending on the time scale of fiber orientation). Video analysis was performed with the ImageJ plugin OrientationJ (OJ Dominant Orientation), yielding a dominant direction and coherency of recorded structures for each video frame. The starting time $T_0$ for the orientation process was determined as the time of the frame in which the µfiber solution first covered the whole recording area. For evaluation, the dominant direction difference and coherency difference between consecutive frames was calculated. µFibers were considered oriented when the rolling mean (10 frame width) of the dominant direction difference changed less than 0.1° for the rest of the video and, additionally, the rolling mean (10 frame width) of the coherency difference changed less than 0.1% for at least ten consecutive frames. The according alignment time was calculated as the difference between the time of the frame fulfilling the orientation criteria and $T_0$. Per sample, at least three videos were recorded to determine the mean alignment time.

*Calculation of µFiber Dispersion Concentrations and Production Rates:* µFibers in dispersion are counted with a Neubauer Improved counting chamber (Marienfeld) to determine µfiber concentrations. To calculate possible µfiber production rates per day, the concentration of a µfiber dispersion $c_{\mu\mathrm{fiber}}$ obtained from one fiber spun for a whole working day was multiplied with the volume of the dispersion $V_{\mathrm{disp}}$ to obtain the total µfiber number, and, following, with the number of fibers $n_{\mathrm{f}}$ that can be spun in parallel in the used setup (3 fibers for Setup I, 8 fibers for Setup II). To consider the time needed for fiber sectioning and µfiber purification, the resulting µfiber number is multiplied with the factor 0.8 (based on the assumption that fiber spinning is conducted for four days, followed by one day of sectioning and purification of all fibers produced in the previous four days):

$$\mu\mathit{Fiber\ Production\ Rate} = c_{\mu\text{fiber}} \cdot V_{\text{disp}} \cdot n_{\text{f}} \cdot \frac{0.8}{\text{d}} \qquad (3)$$

*Cell Culture:* L929 mouse fibroblasts were cultured in DMEM containing 10 vol% FBS. For cytotoxicity experiments, cell culture medium was additionally supplemented with 1 vol% AmB; cells of passage 24 were used.

GFP-HUVECs were cultured on fibronectin-coated (20 µg/mL in PBS) tissue culture flasks (75 $cm^2$) in ECGM, supplemented with low-serum growth medium supplement mix and P/S (1 vol%). For vasculogenesis model experiments, ECGM with a 3X supplement mix concentration and 1 vol% P/S, and cells of passages five to ten are used.

MSCs were obtained from bone marrow aspirates of healthy donors as reported elsewhere.[6] Informed written consent was given by the donors; the study was approved by the local ethics committee (ethical approval no. 307082018). MSCs were cultured in DMEM, supplemented with 10 vol% FBS and 1 vol% P/S. Cells of passages two to five are used. For experiments, MSCs encapsulated together with HUVECs in a hydrogel were cultured using 3X ECGM.

All cell cultures were maintained at 37 °C, 5% $CO_2$ and 90% humidity.

*µFiber Disinfection for Cell Culture Experiments:* Aqueous µfiber dispersions were disinfected using UV irradiation for 3 h under continuous stirring.

*µFiber Cytotoxicity Evaluation with Mouse Fibroblasts*: L929 mouse fibroblasts were seeded in a 96 well plate (12,500 cells/well) and incubated (37°C, 5% $CO_2$, 90% humidity) for 24 h to allow cell attachment. 1 – 3 µL of highly concentrated µfiber dispersion (20,000 µfibers/well) were added into the wells and the well plate was incubated for further 24 h. To wash the µfibers away, the wells were washed with fresh medium, gently pipetted up and down three times per well. A live/dead assay was conducted according to the manufacturer’s protocol. Live (cells cultured as described, without µfibers) and dead (cells cultured as described, without µfibers, treated with EtOH before staining) controls were conducted in parallel. Brightfield and fluorescent images were acquired with a spinning disk confocal fluorescence microscope (Opera Phenix™ Plus high-content imaging system with Harmony software, Revvity) with 488 and 568 nm laser excitation wavelength, cell viability was evaluated by automated quantification of alive and dead cells. Per control or µfiber sample, four or five replicates were analyzed.

*Preparation of High-Throughput 3D Vasculogenesis Model*: Based on vasculogenesis models established by Dennison and Fusenig *et al.*,[1] (starPEG-sGAG)-based hydrogels with a solid content of 1.85 wt% (0.50 mM sGAG, 0.75 mM starPEG-peptide conjugate), containing cell-adhesive RGDSP peptide (1 mM, 2 mol/mol sGAG) and growth factors VEGF, FGF2 and SDF1α (20 µg/mL each) were prepared without or with µfibers ($5.6 \pm 1.2$ µm diameter, $51 \pm 3.3$ µm long; $18 \cdot 10^6$ fibers/mL in µfiber stock dispersion, equals 0.56 vol% µfiber occupancy in hydrogel).

Two hydrogel precursor solutions, one containing starPEG-peptide conjugate, the other containing sGAG, RGDSP peptide, growth factors VEGF, FGF2 and SDF1α, and cell suspensions of HUVECs and MSCs in PBS (termed sGAG precursor solution in the following), were prepared freshly in PBS on the experiment day. The gelation time of the hydrogel forming upon mixing of the two precursor solutions was adjusted to $60 - 80$ s through acidification of the starPEG precursor solution with hydrochloric acid; prior to mixing both precursor solutions together, the starPEG solution used for the respective gelation test was diluted 1:1 with µfiber dispersion (in PBS) or PBS, to test the gelation time with or without µfibers.

Two 96 PCR well plates were filled manually with the precursor solutions: in plate 1, 3 µL of starPEG solution and 3 µL of µfiber dispersion or PBS per well were added. sGAG precursor solution was transferred into plate 2 (8 µL/well). Both plates, together with a 96 µwell plate (µ-Plate 96 Well Round, Ibidi) serving as experiment well plate, were transferred into a liquid handling station (JANUS® G3, part of Explorer™ G3 workstation, Revvity). Further steps were conducted in an automated manner with the liquid handling station (Explorer™ G3 workstation, Revvity), using a 96 channel multi-dispensing unit to handle all targeted wells in parallel: the sGAG precursor solution in plate 2 was mixed to resuspend the cells in solution; 6 µL of the solution were aspirated, transferred to plate 1, and mixed with the respective precursor solution in plate 1. 10 µL of the resulting hydrogel solution were transferred into the 96 µwell experiment plate. Previous adjustment of the gelation time enabled the transfer into the experiment plate while the hydrogel mixture was still pipettable. To allow complete gelation, medium (50 µL/well) was added onto the gels after a latency time of 10 min.

Preparation of hydrogels without or with randomly oriented µfibers was conducted as described above. For the production of hydrogels featuring aligned µfibers, a custom-made magnetic lid suitable for 96 well plates, creating a weak magnetic field ($50 \pm 10$ mT) in every second row of the well plate (Figure S21a), was mounted onto the experiment well plate. The steps performed with the liquid handling workstation were repeated, using the same precursor and experiment plate as before. This was rendered possible as pipet tips, in the procedure described above, were only mounted onto the 96 channel multi-dispensing unit in the positions treating the wells targeted for hydrogel cultures without or with randomly oriented µfibers, leaving the wells in precursor plate 1 and 2 intended for aligned µfiber cultures filled for the repetition, and enabling the preparation of randomly oriented and aligned µfiber cultures in one experiment plate. The custom-made magnetic well plate lid was removed from the experiment plate before medium addition.

Per condition, three to four hydrogel replicates were cultured. Medium change and confocal fluorescence imaging (5x objective, brightfield and 488 nm excitation wavelength, z-stack with 371 µm height and 26.5 µm plane distance) were performed every day and every other day, in an automated manner (using JANUS® G3 liquid handling system and plate::handler™

Flex robotic arm for medium change and Opera Phenix™ Plus high-content imaging system for imaging, all modules part of Explorer™ G3 workstation, Revvity).

*Alignment Analysis of HUVEC Structures:* Raw images of z-stacks (106 µm, five planes, middle portion of originally acquired z-stacks) were processed with ImageJ to obtain one maximum intensity projection per analyzed well. Image analysis was conducted with a customized python code (https://github.com/tomio13/ImageP/blob/main/Example/Gauss-edge.py) as described previously.[7] Briefly, images were background-corrected by subtracting the image resulting from convolution of a normalized Gaussian kernel (10 px width, 61 px window size). Resulting images were smoothed using a convolution filter employing again a normalized Gaussian kernel (1 px width, 11 px window size). Blob suppression was performed by masking the results (in the angle image described below) based on a structure tensor (5 px width, 51 px window size) analysis of the background filtered image to remove pixels with a correlation smaller than 0.5. To detect the orientation angle of imaged structures, an elliptic Mexican hat filter (0.5 px width in one direction and 10 px width perpendicular to it, window size of 21 px), covering 20 angle values between 0 and 180°, was applied. Resulting images were collected in a z-stack with orientation angles varying along the z-axis. A maximum projection along the z-axis was used to identify local maxima and the corresponding orientation angle for each pixel (forming the 'angle image'). The maximum image was filtered using Otsu's threshold to validate pixels and background, keeping only the object pixels for further analysis. Orientation histograms were calculated from the corresponding angle values. Resulting full widths at half maxima (FWHM) of these histograms were employed as a measure of HUVEC structure alignment.

*Statistical analysis:* Statistical analysis of obtained data was performed with OriginPro 2024b using ANOVA with post-hoc Bonferroni test, yielding $p$-values of pairwise comparisons.

$p$-Values below 0.05 were considered to indicate a significant difference ($*p < 0.05$; $**p < 0.01$; $***p < 0.001$; $****p < 0.0001$).

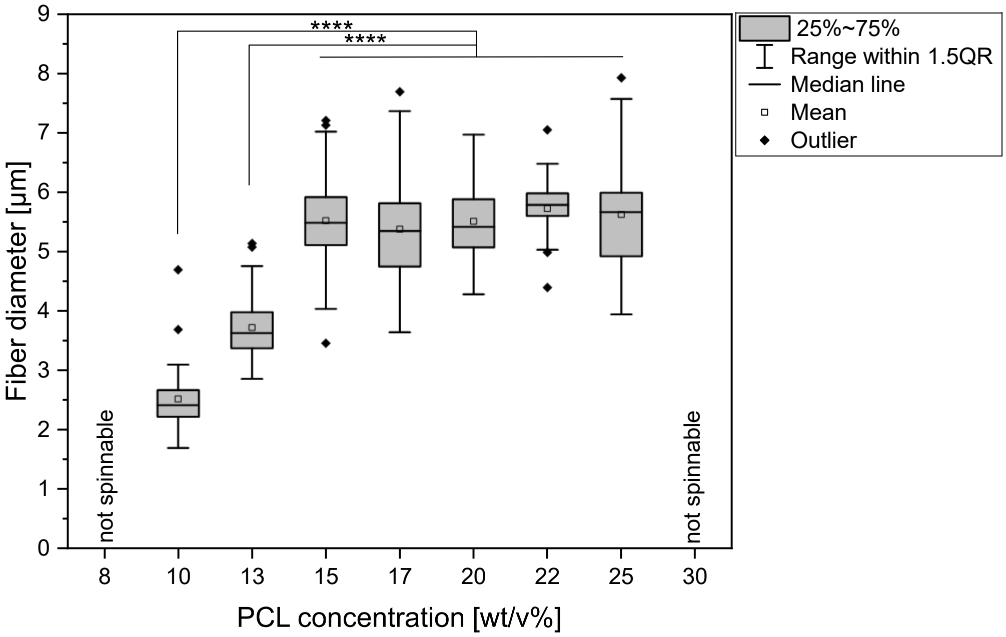


Figure S1: Fiber diameters ($n \geq 40$ per sample) in relation to PCL concentration of the spinning solution. All fibers spun under otherwise same spinning conditions (Setup II, flow rate: 0.3 mL/h, drum speed: 78.5 cm/s); $****p < 0.0001$ (ANOVA, post-hoc Bonferroni test).

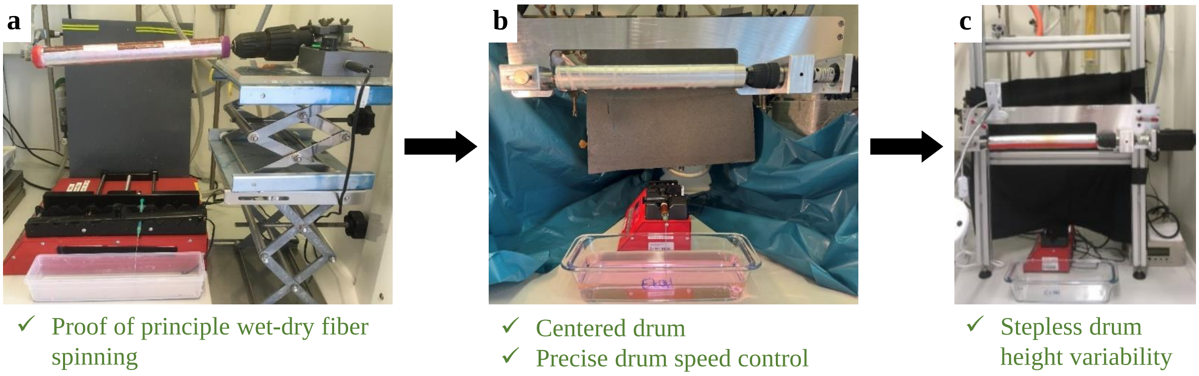


Figure S2: Development of wet-dry spinning setup, resulting in (c) Setup I used for DoE I in this study.

**a**

| | Input Factor | Levels (Coding) | | | | |
|---|---|---|---|---|---|---|
| **1** | Flow rate [mL/h] | 0.0500 (-2) | 0.1125 (-1) | 0.1750 (0) | 0.2375 (1) | 0.3000 (2) |
| **2** | Drum speed [cm/s] | 7.9 (-2) | 15.7 (-1) | 23.6 (0) | 31.4 (1) | 39.3 (2) |
| **3** | Drum height [cm] | 20 (-2) | 30 (-1) | 40 (0) | 50 (1) | 60 (2) |
| **4** | Horiz. drum - cannula distance [cm] | -10.00 (-2) | -3.75 (-1) | 2.50 (0) | 8.75 (1) | 15.00 (2) |

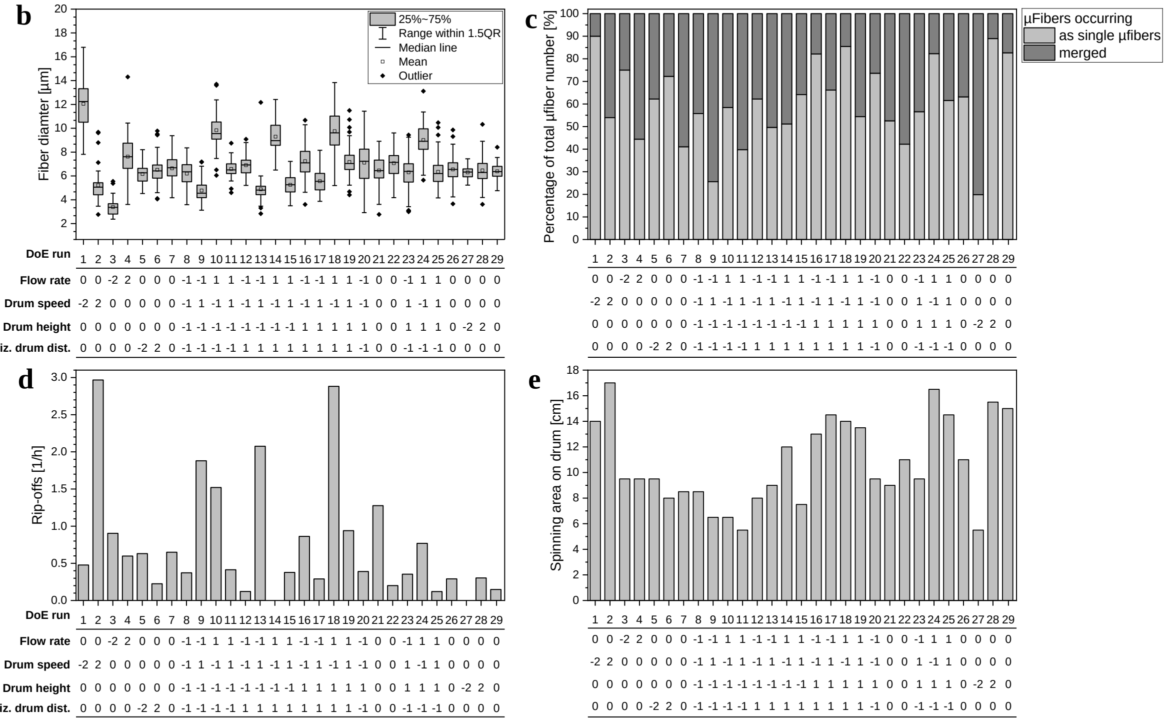


Figure S3: (a) List of input factors and according levels of DoE I (same as Figure 3b, reproduced here for easier readability of plots b – e); blue numbers in brackets give the values normalized to the middle value of the parameter range chosen for the respective input factor, referred to as coding in the following. Experimental results of all DoE I runs: (b) fiber diameters ($n \geq 40$ per sample), (c) amount of µfibers occurring as single µfibers or merged ($n \geq 58$ µfibers counted per sample), (d) average amount of fiber rip-offs occurring per hour, and (e) spinning area of fibers on drum after spinning. Parameter settings of the DoE runs given as coded values introduced in (a).

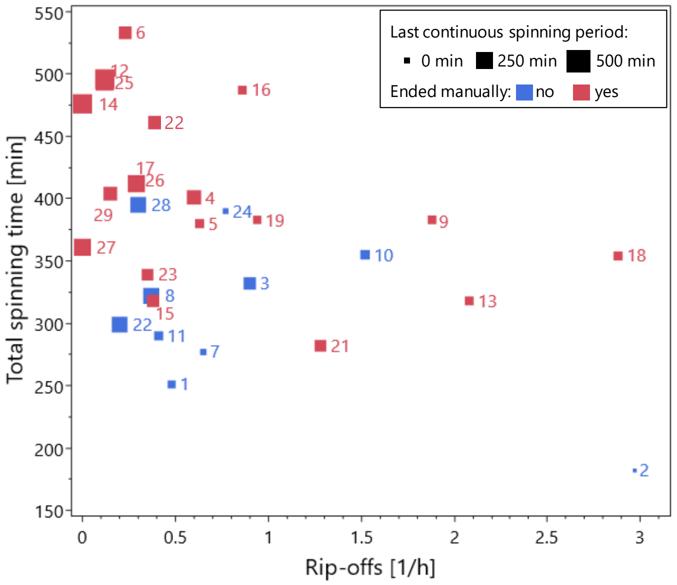


Figure S4: Total spinning times of all DoE I runs plotted against the respective fiber rip-off rates; run numbers are given as numbers next to the data points. Fiber rip-off rates were calculated based on the amount of observed fiber rip-offs during the experiment day and the total spinning time of the experiment. If a fiber was detached manually from the drum at the end of the experiment day, this event was not counted as a fiber rip-off, but the last continuous spinning period was clogged up to this time point and was considered part of the total spinning time. The data point sizes indicate the length of the last continuous spinning period, data points in red denote that this last continuous spinning period of the day was ended manually. It has to be noted that the rip-off rates of these runs, if the last continuous spinning would not have been ended manually, could be slightly different and are thus more prone to deviations than the rate of runs marked in blue. (Last continuous spinning periods ended manually were, nonetheless, not excluded from the analysis because in some runs, no rip-offs were observed during the whole day, so the run had to be ended manually at the end of the experiment day.)

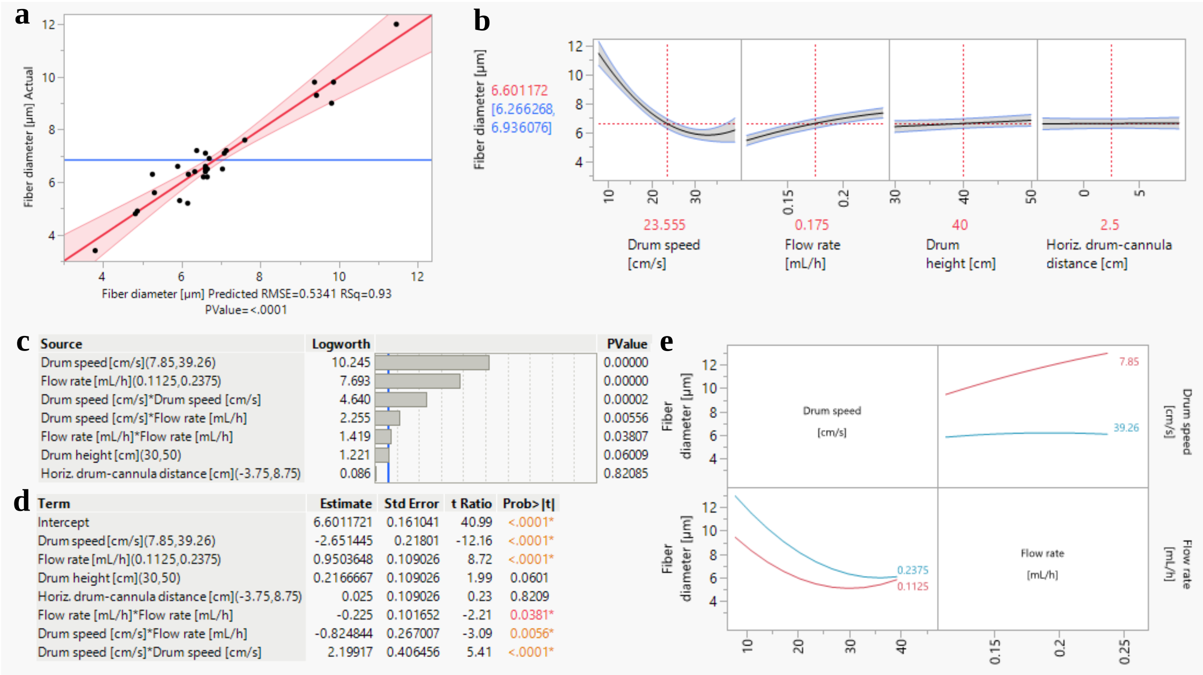


$$6.6011721 - 2.651445 \cdot \left(\frac{Drum\ speed - 23.555}{15.705}\right) + 0.9503648 \cdot \left(\frac{Flow\ rate - 0.175}{0.0625}\right) + 0.2166667 \cdot \left(\frac{Drum\ height - 40}{10}\right) + 0.025 \cdot \left(\frac{Horiz.d.c.dist. - 2.5}{6.25}\right)$$
$$- 0.824844 \cdot \left(\frac{Drum\ speed - 23.555}{15.705}\right) \cdot \left(\left(\frac{Flow\ rate - 0.175}{0.0625}\right)\right) + 2.19917 \cdot \left(\frac{Drum\ speed - 23.555}{15.705}\right)^2 - 0.225 \cdot \left(\frac{Flow\ rate - 0.175}{0.0625}\right)^2$$

Figure S5: DoE I correlative model of the effects of investigated process parameters (flow rate, drum speed, drum height, horizontal drum – cannula distance) on **fiber diameter**, obtained by regression analysis of the experimental DoE data; plots created with JMP software. (a) Whole-model leverage plot, showing obtained experimental data plotted against the predicted fiber diameter at corresponding parameter settings (black dots), ideal model where $y = x$ (red line) with confidence intervals for a significance level of $\alpha = 0.05$ (red areas), and experimental response mean (blue line). As evaluation metrics, root mean square error (RMSE), $R^2$ (RSq) and $p$-value of ANOVA are displayed below the leverage plot. (b) Prediction profilers allowing to interactively predict the fiber diameter for any input factor combination within the tested parameter space and visualizing the main and interaction effects of the input factors. (c) Effect summary plot displaying the $-\log$ transformations of the $p$-value of the main and interaction effects included in the model, sorted from most influential effect at the top to least influential one at the bottom; the blue line indicates the significance level for $\alpha = 0.05$. Main effects, i.e. linear effects of the input factors, are kept in the model even if not significant while non-significant quadratic or non-linear interaction effects were stepwise removed from the model. (d) Factor estimates, corresponding standard errors, t ratios (ratio of estimate to standard error) and $p$-values of the effects included in the model. The estimates are the model coefficients which are used to calculate the predicted fiber diameters; the algebraic sign in front indicates whether the effects have an in- or decreasing influence. (e) Interaction plots of non-linear interaction between drum speed and flow rate, displaying the varying influence of one input factor on the fiber diameter while the other one is set at a higher or lower level. (f) Prediction equation based on the estimates shown in (d).

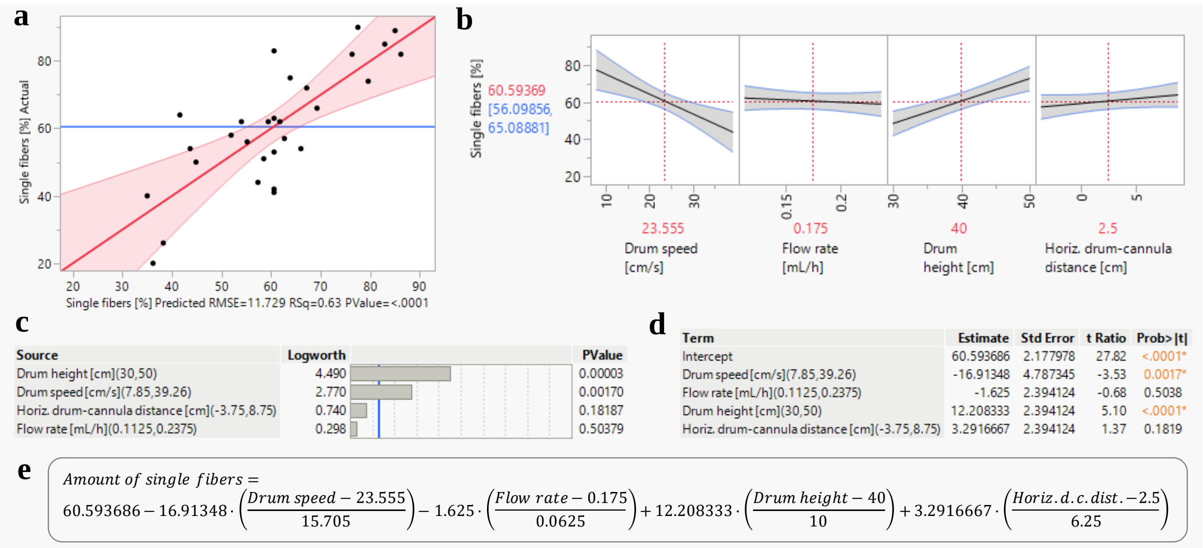


Figure S6: DoE I correlative model of the effects of investigated process parameters (flow rate, drum speed, drum height, horizontal drum – cannula distance) on **µfiber merging**, obtained by regression analysis of the experimental DoE data; plots created with JMP software. (a) Whole-model leverage plot for amount of µfibers occurring as singe fibers, showing obtained experimental data plotted against the predicted data at corresponding parameter settings (black dots), the ideal model where $y = x$ (red line) with a confidence interval for a significance level of $\alpha = 0.05$ (red area), and experimental response mean (blue line). As evaluation metrics, root mean square error (RMSE), $R^2$ (RSq) and $p$-value of ANOVA are displayed below the leverage plot. (b) Prediction profilers allowing to interactively predict the amount of single, non-merged µfibers for any input factor combination within the tested parameter space and visualizing the main effects of the input factors. (c) Effect summary plot displaying the $-\log$ transformations of the $p$-value of the main effects included in the model, sorted from most influential effect at the top to least influential one at the bottom; the blue line indicates the significance level for $\alpha = 0.05$; all interaction effects were not significant and, hence, removed from the model. (d) Factor estimates, corresponding standard errors, t ratios (ratio of estimate to standard error) and $p$-values of the effects included in the model. The estimates are the model coefficients which are used to calculate the predicted amount of µfibers (in percent) occurring as single, non-merged, µfibers; the algebraic sign in front indicates whether the effects have an in- or decreasing influence on the amount of µfibers occurring singly. (e) Prediction equation based on the estimates shown in (d). As no non-linear interaction of two different input factors is contained in the model, no interaction plots are displayed here.

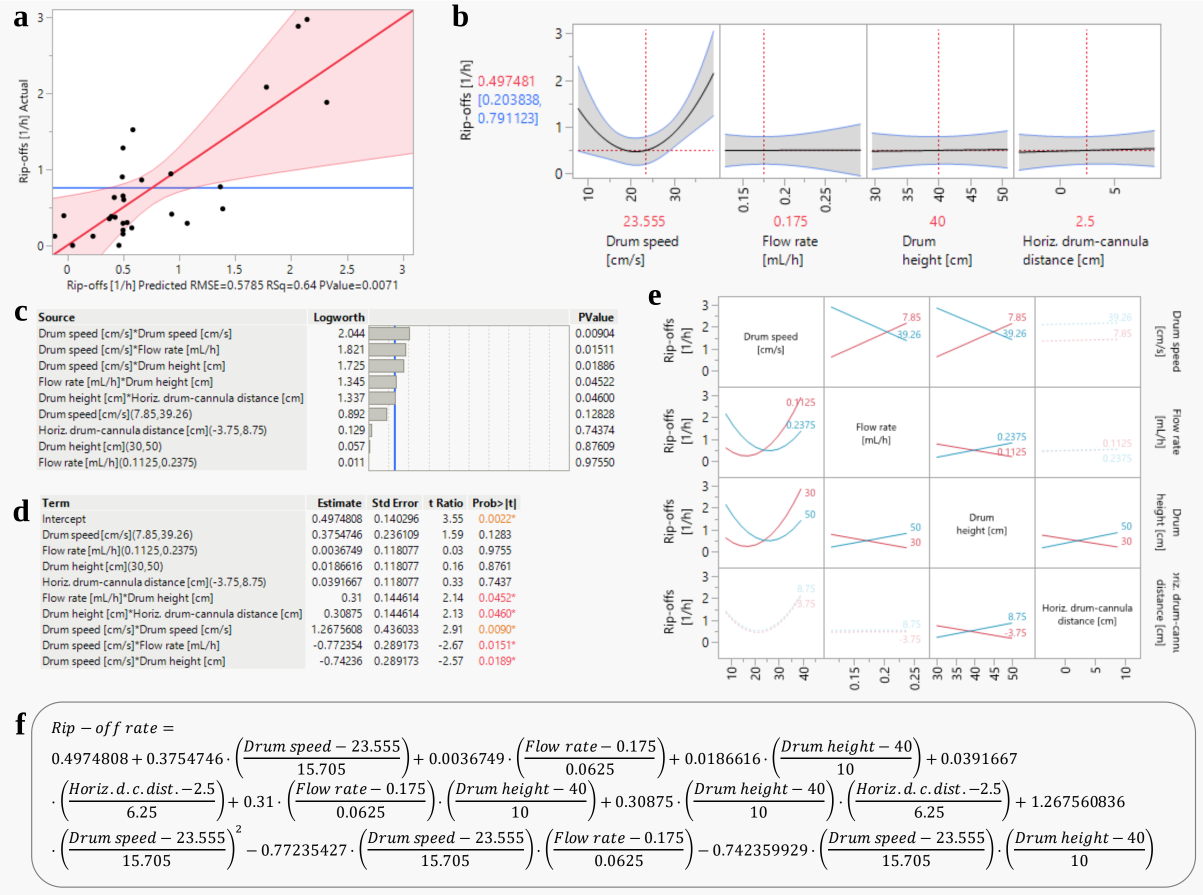


Figure S7: DoE I correlative model of the effects of investigated process parameters (flow rate, drum speed, drum height, horizontal drum – cannula distance) on the average amount of **fiber rip-offs** observed during a respective run, obtained by regression analysis of the experimental DoE data; plots created with JMP software. (a) Whole-model leverage plot, showing obtained experimental data plotted against the predicted rip-off rate at corresponding parameter settings (black dots), ideal model where $y = x$ (red line) with a confidence interval for a significance level of $\alpha = 0.05$ (red area), and experimental response mean (blue line). As evaluation metrics, root mean square error (RMSE), $R^2$ (RSq) and $p$-value of ANOVA are displayed below the leverage plot. (b) Prediction profilers allowing to interactively predict the average amount of rip-offs for any input factor combination within the tested parameter space and visualizing the main and interaction effects of the input factors. (c) Effect summary plot displaying the $-\log$ transformations of the $p$-value of the main and interaction effects included in the model, sorted from most influential effect at the top to least influential one at the bottom; the blue line indicates the significance level for $\alpha = 0.05$. Main effects, i.e. linear effects of the input factors, are kept in the model even if not significant while non-significant quadratic or non-linear interaction effects were stepwise removed from the model. (d) Factor estimates, corresponding standard errors, t ratios (ratio of estimate to standard error) and *p*-values of the effects included in the model. The estimates are the model coefficients which are used to calculate the predicted average amount of rip-offs per hour; the algebraic sign in front indicates whether the effects have an in- or decreasing influence. (e) Interaction plots of non-linear interactions between drum speed, flow rate, drum height and horizontal drum distance, displaying the varying influence of one input factor on continuous spinning while the other one is set at a higher or lower level. (f) Prediction equation based on the estimates shown in (d).

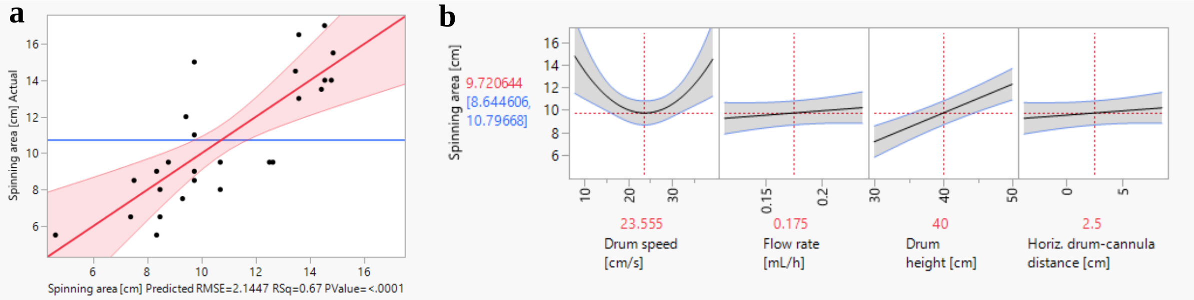


**c**

| Source | Logworth | PValue |
|---|---|---|
| Drum height [cm](30,50) | 5.239 | 0.00001 |
| Drum speed [cm/s]*Drum speed [cm/s] | 2.246 | 0.00567 |
| Flow rate [mL/h](0.1125,0.2375) | 0.545 | 0.28505 |
| Horiz. drum-cannula distance [cm](-3.75,8.75) | 0.545 | 0.28505 |
| Drum speed [cm/s](7.85,39.26) | 0.054 | 0.88383 |

**d**

| Term | Estimate | Std Error | t Ratio | Prob>\|t\| |
|---|---|---|---|---|
| Intercept | 9.7206445 | 0.520163 | 18.69 | <.0001* |
| Drum speed [cm/s](7.85,39.26) | -0.129338 | 0.8754 | -0.15 | 0.8838 |
| Flow rate [mL/h](0.1125,0.2375) | 0.4791667 | 0.437782 | 1.09 | 0.2851 |
| Drum height [cm](30,50) | 2.5625 | 0.437782 | 5.85 | <.0001* |
| Horiz. drum-cannula distance [cm](-3.75,8.75) | 0.4791667 | 0.437782 | 1.09 | 0.2851 |
| Drum speed [cm/s]*Drum speed [cm/s] | 4.9319622 | 1.616642 | 3.05 | 0.0057* |

**e**

$$Spinning\ area = 9.7206445 - 0.129338 \cdot \left(\frac{Drum\ speed - 23.555}{15.705}\right) + 0.4791667 \cdot \left(\frac{Flow\ rate - 0.175}{0.0625}\right) + 2.5625 \cdot \left(\frac{Drum\ height - 40}{10}\right) + 0.4791667 \cdot \left(\frac{Horiz.d.-c.dist.-2.5}{6.25}\right) + 4.9319622 \cdot \left(\frac{Drum\ speed - 23.555}{15.705}\right)^2$$

Figure S8: DoE I correlative model of the effects of investigated process parameters (flow rate, drum speed, drum height, horizontal drum – cannula distance) on **spinning area** of fibers on the drum, obtained by regression analysis of the experimental DoE data; plots created with JMP software. (a) Whole-model leverage plot, showing obtained experimental data plotted against the predicted spinning area at corresponding parameter settings (black dots), ideal model where $y = x$ (red line) with a confidence interval for a significance level of $\alpha = 0.05$ (red area), and experimental response mean (blue line). As evaluation metrics, root mean square error (RMSE), $R^2$ (RSq) and *p*-value of ANOVA are displayed below the leverage plot. (b) Prediction profilers allowing to interactively predict the spinning area for any input factor combination within the tested parameter space and visualizing the main and interaction effects of the input factors. (c) Effect summary plot displaying the $-\log$ transformations of the *p*-value of the main and interaction effects included in the model, sorted from most influential effect at the top to least influential one at the bottom; the blue line indicates the significance level for $\alpha = 0.05$. Main effects, i.e. linear effects of the input factors, are kept in the model even if not significant while non-significant quadratic or non-linear interaction effects were stepwise removed from the model. (d) Factor estimates, corresponding standard errors, t ratios (ratio of estimate to standard error) and $p$-values of the effects included in the model. The estimates are the model coefficients which are used to calculate the predicted spinning areas; the algebraic sign in front indicates whether the effects have an in- or decreasing influence. (e) Prediction equation based on the estimates shown in (d). As no non-linear interaction of two different input factors is contained in the model, no interaction plots are displayed here.

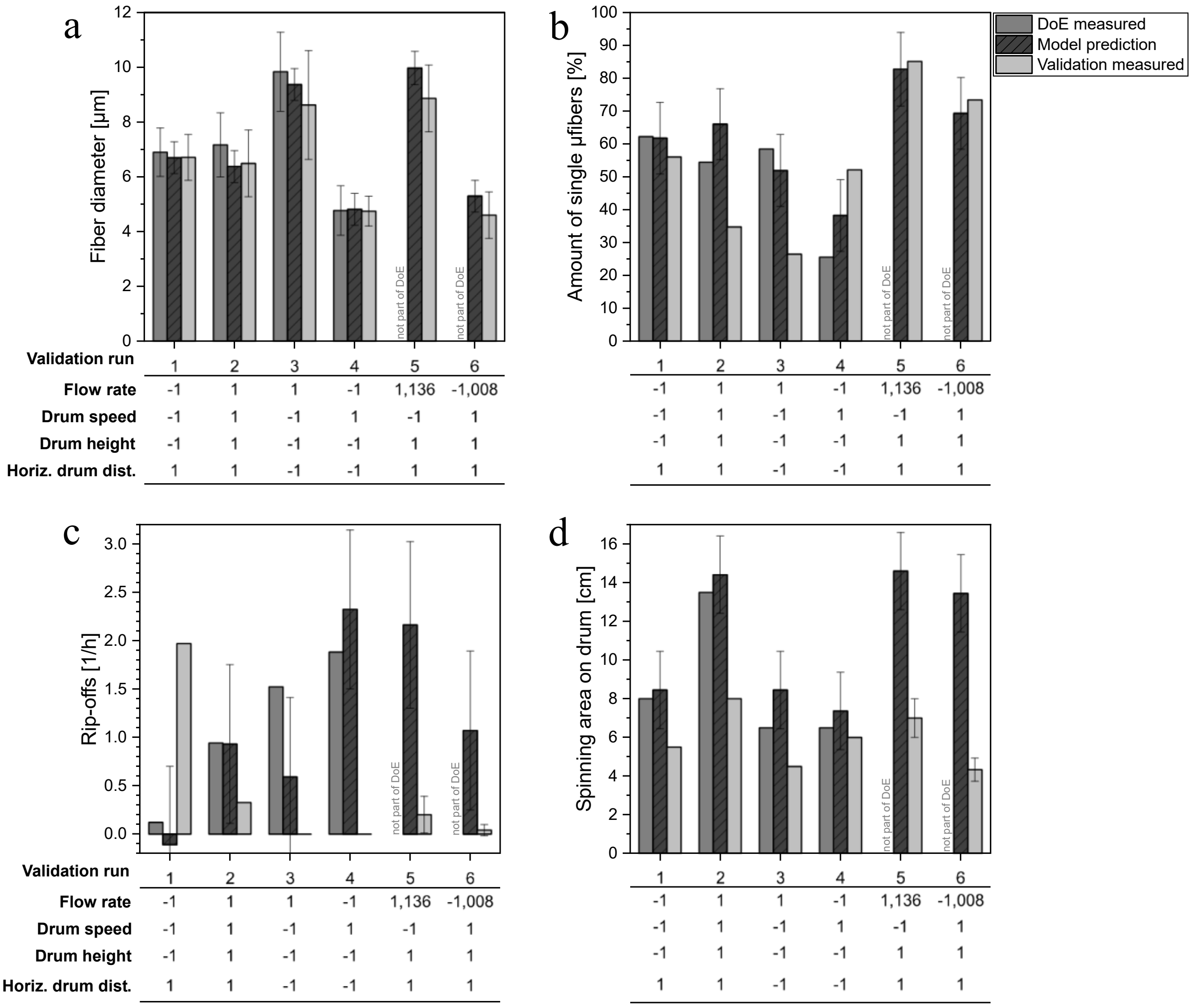


Figure S9: Validation of the statistical model obtained with DoE I. Four runs of the DoE were repeated and two runs with settings that were not part of the DoE were conducted. Results of the original DoE runs, results predicted by the model generated with the DoE and results of the validation runs are shown for (a) fiber diameters ($n \geq 40$ per sample), (b) amount of µfibers occurring as single µfibers ($n \geq 58$ µfibers counted per sample), (c) average amount of fiber rip-offs per hour, and (d) spinning area of fibers on drum after spinning. Error bars of the model predictions indicate the respective 95% confidence interval; error bars of DoE and validation run results indicate the standard deviation of respective mean values if $n > 1$ (i.e. in case of fiber diameters and for validation runs 5 and 6 in (c) and (d), as three fibers were spun in parallel here; for other DoE and validation runs only one fiber was spun per run).

*Analysis of DoE II data:* A full factorial design was chosen to screen the output parameters at extreme spinning conditions, varying flow rate and drum speed as input factors. As a two factor – full factorial design only allows to detect linear main effects ($X_1, X_2$) and the non-linear interaction effect ($X_1 X_2$), but no quadratic effects of the factors ($X_1^2, X_2^2$), a center point was included in the design to permit quadratic terms during fitting. In this design, the quadratic effects are aliased due to a limited amount of degrees of freedom during fitting, thus, it can be analyzed if quadratic effects exist but not which of the effects specifically contributes.[8] Nonetheless, this design approach is commonly used for screening and sufficient to observe how the output parameters are effected towards the chosen boundaries. Here, the quadratic interaction effect of the flow rate was utilized for fitting as suggested by JMP software; the quadratic interaction effect of the drum speed was therefore not included.

Due to the spinnability window and the resulting adjustment of the design point $0.05\ \mathrm{mL/h}\ (-1)$ / $523.6\ \mathrm{cm/s}\ (1)$ to $0.05\ \mathrm{mL/h}\ (-1)$ / $144.0\ \mathrm{cm/s}\ (-0.71)$, the DoE II design was not symmetric, restricting the power of the DoE approach. Hence, fitted models here, in contrast to DoE I, should not be used to precisely predict output parameters throughout the parameter space but can be used to identify trends between the obtained experimental data at extreme spinning conditions. For precise prediction within the DoE II parameter space, areas of interest could be investigated by adding further points to the design in future studies to increase the power of the design.

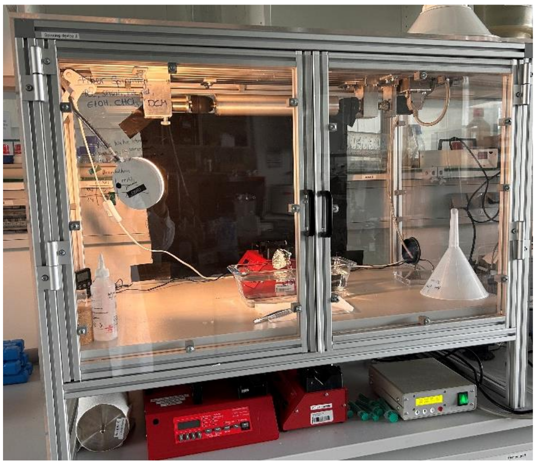

- ✓ Shielding against air movement
- ✓ Higher drum speeds up to 1047 cm/s
- Drum height: 40 cm
- Horiz. drum dist.: -7 cm

Figure S10: Spinning Setup II. The rotating drum collector is built into an acrylic glass box to shield fibers during spinning from environmental air movement, and a motor enabling rotational drum speeds up to 4000 rpm, resulting in surface speeds of up to 1047 cm/s with a drum diameter of 5 cm, is implemented. The drum height and horizontal drum – cannula distance are fixed to 40 cm and −7 cm (cannula in side view left of clockwise rotating drum, as shown in Figure 3d), respectively.

**a**

| | Input Factor | Levels (Coding) | | | |
|---|---|---|---|---|---|
| 1 | Flow rate [mL/h] | 0.050 (-1) | | 0.175 (0) | 0.300 (1) |
| 2 | Drum speed [cm/s] | 78.5 (-1) | 144.0 (-0.71) | 301.1 (0) | 523.6 (1) |

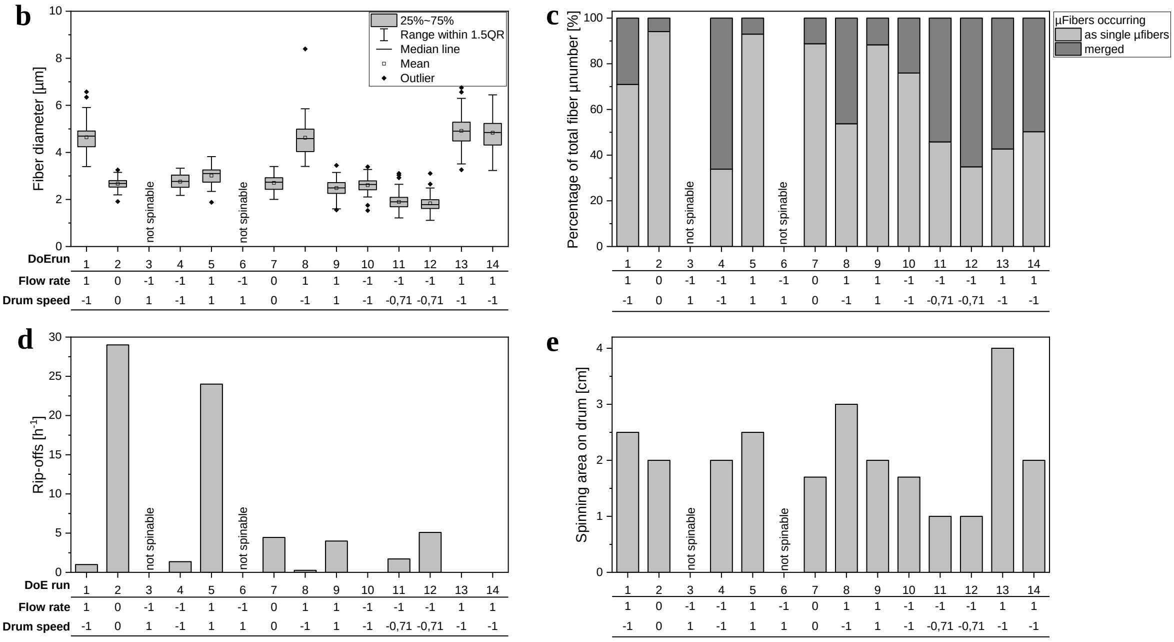


Figure S11: (a) List of input factors and according levels of DoE II; blue numbers in brackets give the values normalized to the middle value of the parameter range chosen for the respective input factor, referred to as coding in the following. Experimental results of all DoE II runs: (b) fiber diameters ($n \geq 40$ per sample), (c) amount of µfibers occurring as single µfibers or merged ($n \geq 58$ µfibers counted per sample), (d) average amount of fiber rip-offs occurring per hour, and (e) spinning area of fibers on drum after spinning. Parameter settings of the DoE runs given as coded values as introduced in (a).

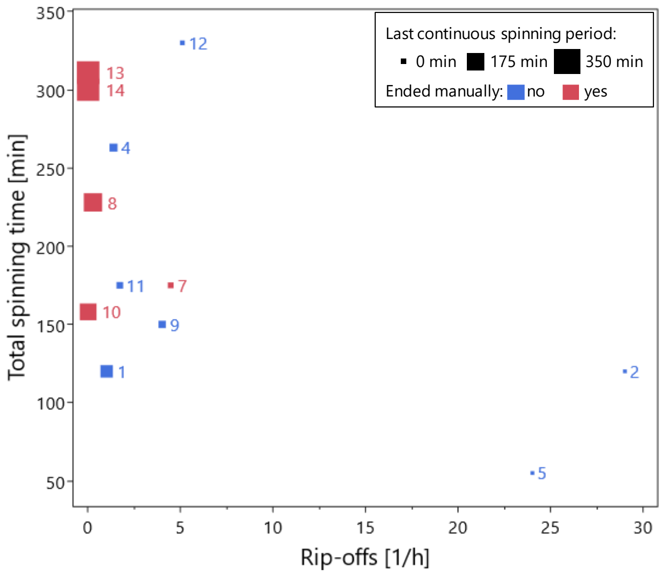


Figure S12: Total spinning times of all DoE II runs plotted against the respective fiber rip-off rates; run numbers are given as numbers next to the data points. Fiber rip-off rates were calculated based on the amount of observed fiber rip-offs during the experiment day and the total spinning time of the experiment. If a fiber was detached manually from the drum at the end of the experiment, this event was not counted as a fiber rip-off, but the last continuous spinning period was clogged up to this time point and was considered part of the total spinning time. The data point sizes indicate the length of the last continuous spinning period, data points in red denote that this last continuous spinning period of the day was ended manually. It has to be noted that the rip-off rates of these runs, if the last continuous spinning would not have been ended manually, could be slightly different and are thus more prone to deviations than the rate of runs marked in blue. (Last continuous spinning periods ended manually were, nonetheless, not excluded from the analysis because in some runs, no rip-offs were observed during the whole experiment, so the run had to be ended manually.)

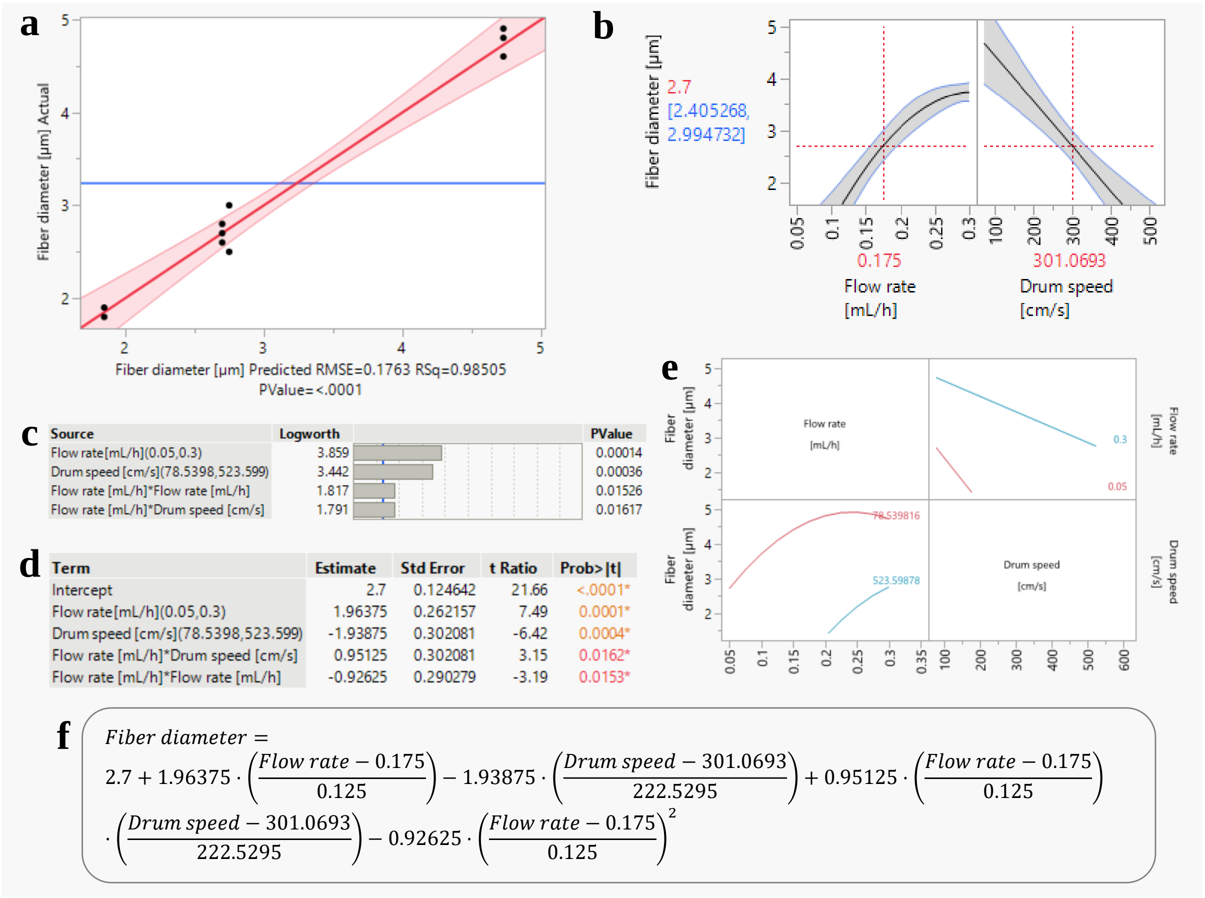


Figure S13: DoE II correlative model of the effects of flow rate and drum speed on **fiber diameter**, obtained by regression analysis of the experimental DoE data; plots created with JMP software. (a) Whole-model leverage plot, showing obtained experimental data plotted against the predicted fiber diameter at corresponding parameter settings (black dots), ideal model where $y = x$ (red line) with confidence intervals for a significance level of $\alpha = 0.05$ (red areas), and experimental response mean (blue line). As evaluation metrics, root mean square error (RMSE), $R^2$ (RSq) and $p$-value of ANOVA are displayed below the leverage plot. (b) Prediction profilers allowing to interactively predict the fiber diameter for any input factor combination within the tested parameter space and visualizing the main and interaction effects of the input factors. (c) Effect summary plot displaying the $-$log transformations of the $p$-value of the main and interaction effects included in the model; the blue line indicates the significance level for $\alpha = 0.05$. All main and interaction effects used for initial fitting are significant, therefore, no effect was removed from the model. (d) Factor estimates, corresponding standard errors, t ratios (ratio of estimate to standard error) and $p$-values of the effects included in the model; the algebraic sign in front of the estimates indicates whether the effects have an in- or decreasing influence. (e) Interaction plots of non-linear interaction between drum speed and flow rate, displaying the varying influence of one input factor on the fiber diameter while the other one is set at a higher or lower level. (f) Prediction equation based on the estimates shown in (d).

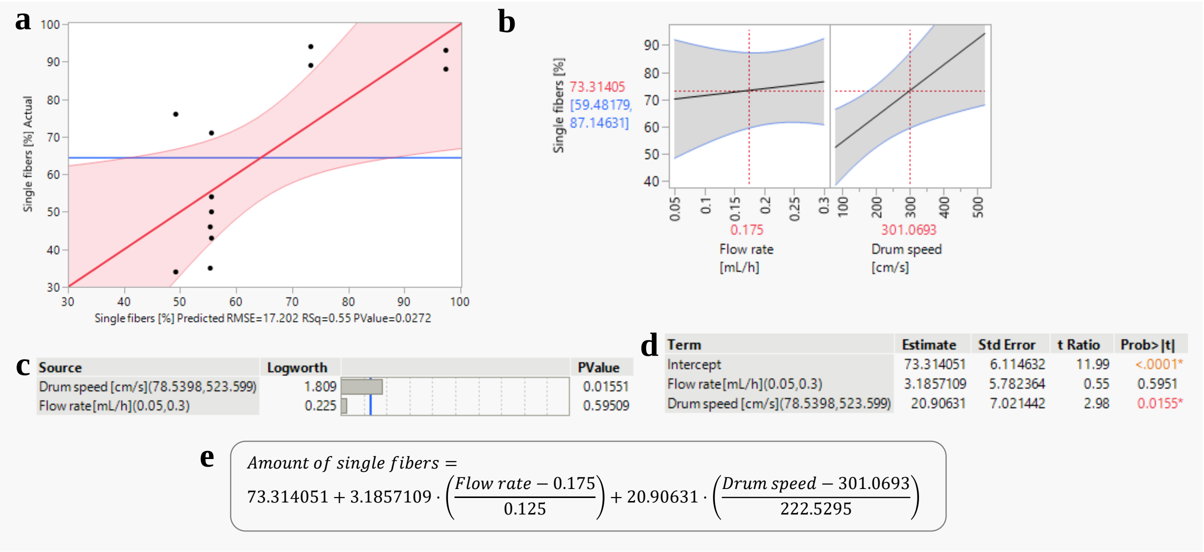


Figure S14: DoE II correlative model of the effects of flow rate and drum speed on **µfiber merging**, obtained by regression analysis of the experimental DoE data; plots created with JMP software. (a) Whole-model leverage plot for µfibers occurring as single fibers in a µfiber dispersion, showing obtained experimental data plotted against the predicted data at corresponding parameter settings (black dots), the ideal model where $y = x$ (red line) with a confidence interval for a significance level of $\alpha = 0.05$ (red area), and experimental response mean (blue line). As evaluation metrics, root mean square error (RMSE), $R^2$ (RSq) and $p$-value of ANOVA are displayed below the leverage plot. (b) Prediction profilers allowing to interactively predict the amount of single, non-merged µfibers for any input factor combination within the tested parameter space and visualizing the main effects of the input factors. (c) Effect summary plot displaying the $-\log$ transformations of the $p$-value of the main effects included in the model. The blue line indicates the significance level for $\alpha = 0.05$.Main effects, i.e. linear effects of the input factors, are kept in the model even if not significant while non-significant quadratic or non-linear interaction effects were stepwise removed from the model. (d) Factor estimates, corresponding standard errors, t ratios (ratio of estimate to standard error) and $p$-values of the effects included in the model; the algebraic sign in front indicates whether the effects have an in- or decreasing influence. (e) Prediction equation based on the estimates shown in (d). As no interaction effects are contained in the model, no interaction plots are displayed here.

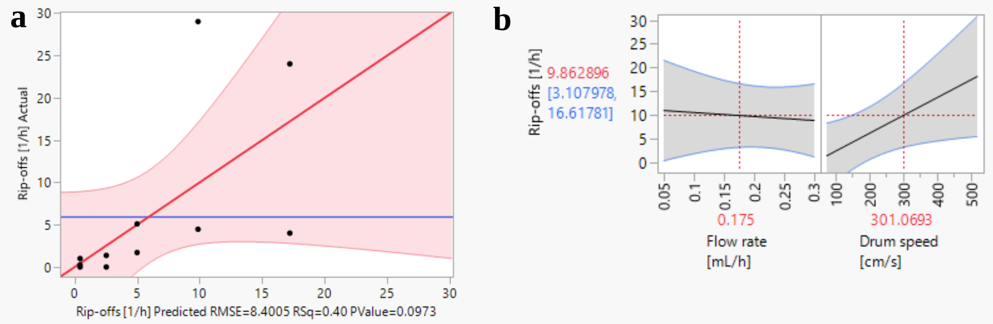


c

| Source | Logworth | PValue |
|---|---|---|
| Drum speed [cm/s](78.5398,523.599) | 1.431 | 0.03706 |
| Flow rate[mL/h](0.05,0.3) | 0.143 | 0.71934 |

d

| Term | Estimate | Std Error | t Ratio | Prob>\|t\| |
|---|---|---|---|---|
| Intercept | 9.8628964 | 2.986052 | 3.30 | 0.0092* |
| Flow rate[mL/h](0.05,0.3) | -1.047156 | 2.82379 | -0.37 | 0.7193 |
| Drum speed [cm/s](78.5398,523.599) | 8.3836691 | 3.428888 | 2.45 | 0.0371* |

e

$$Rip-off\ rate = 9.8628964 - 1.047156 \cdot \left(\frac{Flow\ rate - 0.175}{0.125}\right) + 8.3836691 \cdot \left(\frac{Drum\ speed - 301.0693}{222.5295}\right)$$

Figure S15: DoE II correlative model of the effects of flow rate and drum speed on the average amount of **fiber rip-offs** observed during a respective run, obtained by regression analysis of the experimental DoE data; plots created with JMP software. (a) Whole-model leverage plot, showing obtained experimental data plotted against the predicted rip-offs at corresponding parameter settings (black dots), ideal model where $y = x$ (red line) with confidence intervals for a significance level of $\alpha = 0.05$ (red areas), and experimental response mean (blue line). As evaluation metrics, root mean square error (RMSE), $R^2$ (RSq) and $p$-value of ANOVA are displayed below the leverage plot. (b) Prediction profilers allowing to interactively predict the average amount of fiber rip-offs per hour for any input factor combination within the tested parameter space and visualizing the main effects of the input factors. (c) Effect summary plot displaying the $-\log$ transformations of the $p$-value of the main effects included in the model; the blue line indicates the significance level for $\alpha = 0.05$. Main effects, i.e. linear effects of the input factors, are kept in the model even if not significant while non-significant quadratic or non-linear interaction effects were stepwise removed from the model. (d) Factor estimates, corresponding standard errors, t ratios (ratio of estimate to standard error) and $p$-values of the effects included in the model; the algebraic sign in front of the estimates indicates whether the effects have an in- or decreasing influence. (e) Prediction equation based on the estimates shown in (d). As no interaction effects are contained in the model, no interaction plots are displayed here.

a b

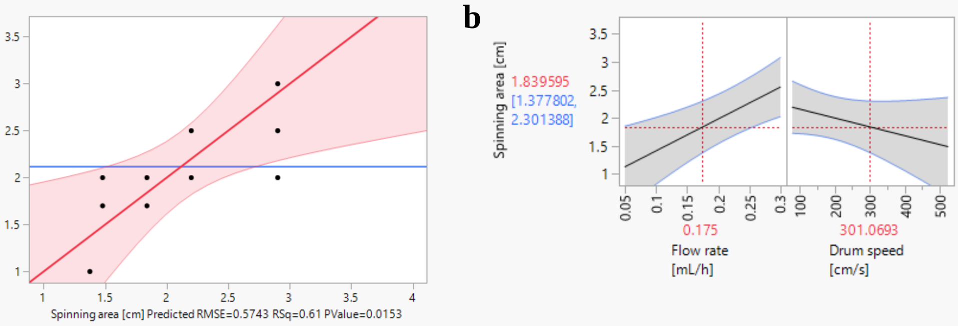


**c**

| Source | Logworth | PValue |
|---|---|---|
| Flow rate[mL/h](0.05,0.3) | 2.299 | 0.00502 |
| Drum speed [cm/s](78.5398,523.599) | 0.774 | 0.16813 |

**d**

| Term | Estimate | Std Error | t Ratio | Prob>\|t\| |
|---|---|---|---|---|
| Intercept | 1.8395953 | 0.204138 | 9.01 | <.0001* |
| Flow rate[mL/h](0.05,0.3) | 0.711685 | 0.193045 | 3.69 | 0.0050* |
| Drum speed [cm/s](78.5398,523.599) | -0.351362 | 0.234412 | -1.50 | 0.1681 |

**e**

$$\mathit{Spinning\ area} = 1.8395953 + 0.711685 \cdot \left(\frac{\mathit{Flow\ rate} - 0.175}{0.125}\right) - 0.351362 \cdot \left(\frac{\mathit{Drum\ speed} - 301.0693}{222.5295}\right)$$

Figure S16: DoE II correlative model of the effects of flow rate and drum speed on the **spinning area** of fibers on the drum, obtained by regression analysis of the experimental DoE data; plots created with JMP software. (a) Whole-model leverage plot, showing obtained experimental data plotted against the predicted spinning area at corresponding parameter settings (black dots), ideal model where $y = x$ (red line) with confidence intervals for a significance level of $\alpha = 0.05$ (red areas), and experimental response mean (blue line). As evaluation metrics, root mean square error (RMSE), $R^2$ (RSq) and $p$-value of ANOVA are displayed below the leverage plot. (b) Prediction profilers allowing to interactively predict the spinning area for any input factor combination within the tested parameter space and visualizing the main and interaction effects of the input factors. (c) Effect summary plot displaying the $-\log$ transformations of the $p$-value of the main and interaction effects included in the model; the blue line indicates the significance level for $\alpha = 0.05$. Main effects, i.e. linear effects of the input factors, are kept in the model even if not significant while non-significant quadratic or non-linear interaction effects were stepwise removed from the model. (d) Factor estimates, corresponding standard errors, t ratios (ratio of estimate to standard error) and $p$-values of the effects included in the model; the algebraic sign in front of the estimates indicates whether the effects have an in- or decreasing influence. (e) Prediction equation based on the estimates shown in (d). As no interaction effects are contained in the model, no interaction plots are displayed here.

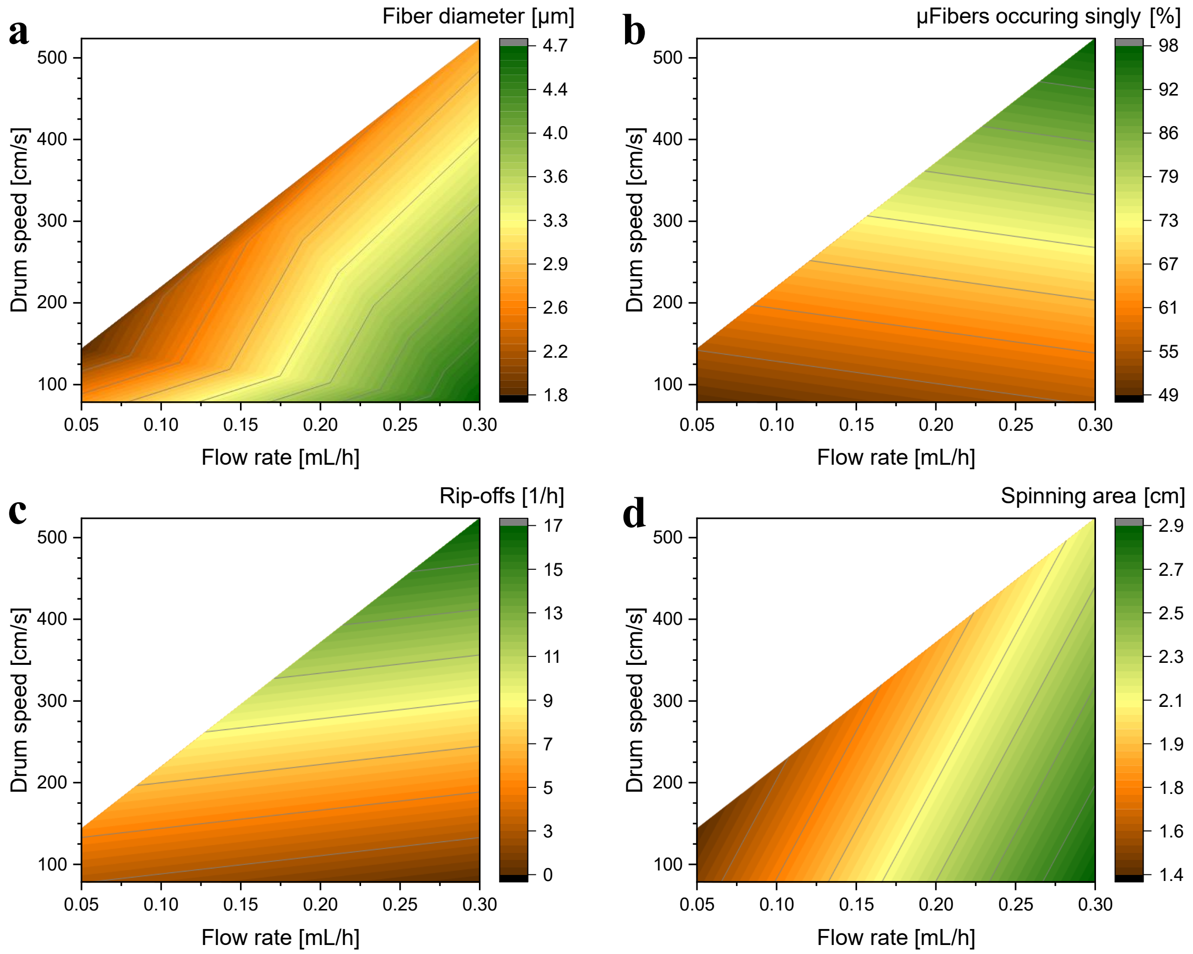


Figure S17: Response surface plots modelled based on prediction equations (Figure S13 – Figure S16) obtained by regression analysis and plotted based on the parameter combinations tested in DoE II, considering the spinnability window. Modelled output parameters: (a) fiber diameter, (b) amount of µfibers occurring individually, not merged, (c) average amount of fiber rip-offs per hour (i.e. rip-off rate), (d) spinning area of fibers on drum after spinning. As noted above, due to restrictions in DoE II, these plots should not be used to precisely predict output parameters throughout the parameter space but can be used to identify trends between the obtained experimental data at extreme spinning conditions.

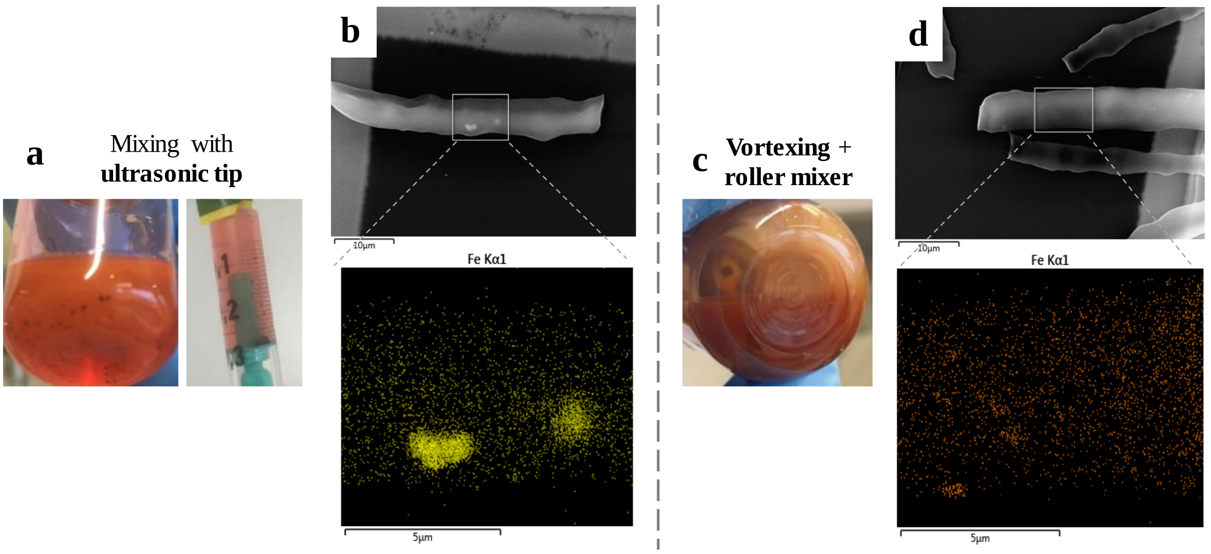


Figure S18: Spinning solutions containing PCL, chloroform, RhB and SPIONs, prepared in two different ways regarding the SPION addition and mixing process. (a) PCL was dissolved in chloroform mixed with RhB stock solution (in chloroform) under continuous stirring for 1 h. SPION stock dispersion (in chloroform) was pipetted onto the viscous solution which was subsequently mixed with an ultrasonic horn tip. (c) Chloroform, RhB stock solution and SPION stock dispersion were mixed by pipetting and vortexing. PCL pellets were added and the solution was vortexed for 1 min and treated on the roller mixer overnight. (b, d) Scanning electron microscopy images (top) and corresponding energy dispersive X-ray spectroscopy images (bottom) showing the iron distribution near the µfiber surface of µfibers produced using spinning solution prepared with (b) ultrasonication or (d) vortexing/roller mixer treatment.

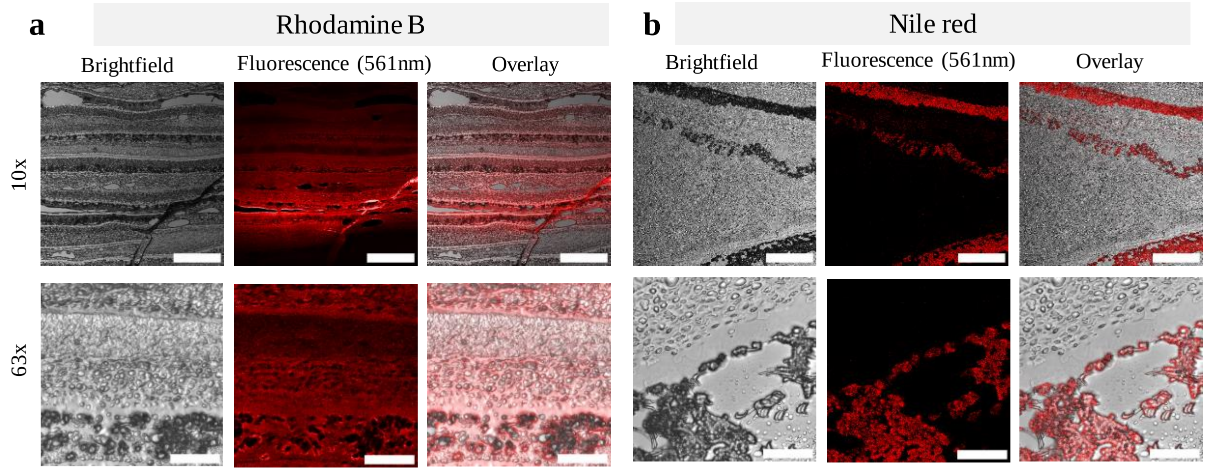


Figure S19: Fluorescence microscopy images of fiber-OCT cross-section slices, collected during cryo-cutting, of fibers with (a) RhB or (b) nile red as dye. Images taken with 10x and 63x objectives. Scale bars: 10x: 300 µm, 63x: 50 µm.

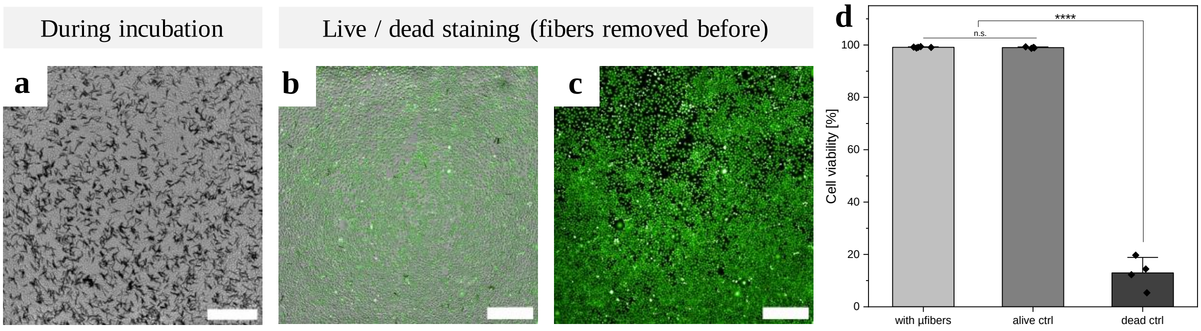


Figure S20: Results of µfiber toxicity test (0.99 wt% SPIONs, 0.01 mg/mL nile red in spinning solution). L929 mouse fibroblasts were incubated one day in direct contact with µfibers. Afterwards, µfibers were removed (because nile red dye in µfibers would interfere with dead dye during imaging) and a live / dead staining was performed. (a) Brightfield microscopy image before live / dead staining, showing attached cells in direct contact with µfibers. (b) Brightfield – fluorescence overlay and (c) fluorescence microscopy image of cells after live / dead staining (excitation with 488 nm and 658 nm, green: alive, red: dead cells). (d) Cell viability of cells incubated with µfibers and live and dead controls. Scale bars: 500 µm.

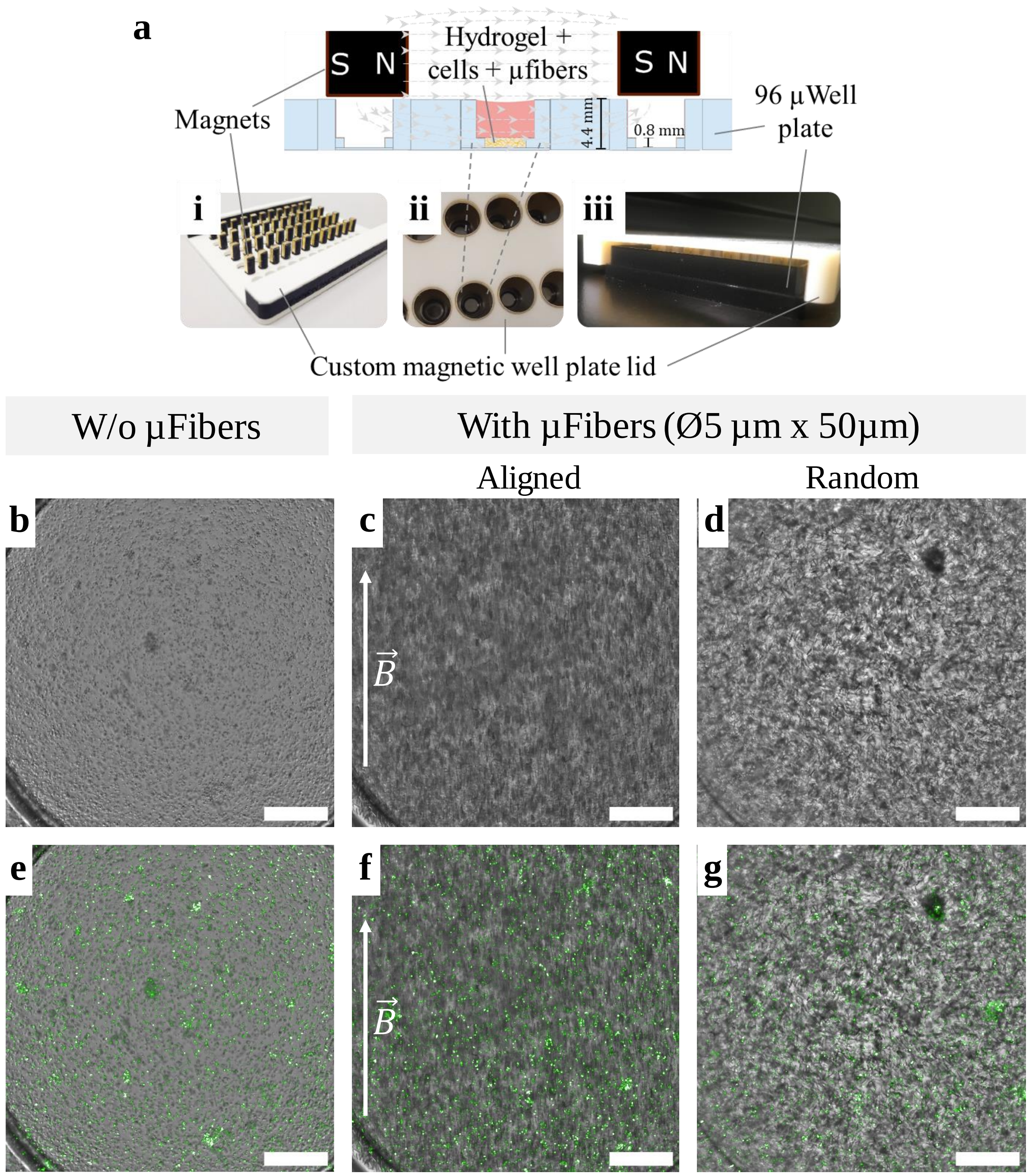


Figure S21: Setup during cell culture model fabrication. (a) A hydrogel containing HUVECs, MSCs and µfibers is casted into a 96 µwell plate equipped with a custom-made magnetic well plate lid. The bottom side of the custom-made lid (i) exhibits magnets in the well positions of every second row of a 96 well plate, while the top plate of the lid (ii) features round openings corresponding to the well openings of the rows between the magnets, through which underlying wells can be filled by pipetting when the magnetic lid is placed on the 96 µwell plate (iii). The magnets create a magnetic field in the wells open for pipetting, enabling the alignment of our µfibers into one direction, along the magnetic field. As the well diameter of the µwell plates used here is smaller than the well diameter of standard 96 well plates, the magnets of the custom-made magnetic lid were mounted on top of the well openings. The magnetic field created by the magnets bridged the 4.4 mm gap between well top and well bottom, creating a sufficiently homogenous magnetic field in the culture well for µfiber alignment. (b – d) Brightfield microscopy images and (e – g) corresponding overlays with confocal fluorescence images of HUVECs (GFP expressing, green) and MSCs cultured in a (starPEG-sGAG)-based hydrogel (b and e) without, (c and f) with magnetically aligned or (d and g) with randomly oriented µfibers at DIV1 (bottom planes of the z-stack projections shown in Figure 7c – e). µFibers used in this experiment were not optimized regarding their fluorescent dye yet and could, therefore, not be visualized with fluorescence microscopy. Nonetheless, their aligned or random orientation is visible in the brightfield images (c and d) here.

Babu, I. Chen, S. Vedaraman, J. Gerardo-Nava, C. Licht, Y. Kittel, T. Haraszti, J. Di Russo, L. De Laporte, *Adv Funct Materials* **2022**, *32*, 2202468.

[8] S. Soravia, A. Orth in *Ullmann's encyclopedia of industrial chemistry* (Eds.: M. Bohnet, F. Ullmann), Wiley-VCH, Weinheim, **2003**.
